# CdrS is a global transcriptional regulator influencing cell division in *Haloferax volcanii*

**DOI:** 10.1101/2021.05.11.443588

**Authors:** Yan Liao, Verena Vogel, Sabine Hauber, Jürgen Bartel, Omer S. Alkhnbashi, Sandra Maaß, Thandi S. Schwarz, Rolf Backofen, Dörte Becher, Iain G. Duggin, Anita Marchfelder

## Abstract

Transcriptional regulators that integrate cellular and environmental signals to control cell division are well known in bacteria and eukaryotes, but their existence is poorly understood in archaea. We identified a conserved gene (*cdrS)* that encodes a small protein and is highly transcribed in the model archaeon *Haloferax volcanii*. The *cdrS* gene could not be deleted, but CRISPRi-mediated repression of the *cdrS* gene caused slow growth, cell division defects, and changed the expression of multiple genes and their products associated with cell division, protein degradation and metabolism. Consistent with this complex regulatory network, overexpression of *cdrS* inhibited cell division, whereas overexpression of the operon encoding both CdrS and a tubulin-like cell division protein (FtsZ2) stimulated division. ChIP-Seq identified 18 DNA-binding sites of the CdrS protein including one upstream of the promoter for diadenylate cyclase, which is an essential gene involved in c-di-AMP signalling implicated in the regulation of cell division. These findings suggest that CdrS is a transcription factor that plays a central role in a regulatory network coordinating metabolism and cell division.

**Importance:** Cell division is a central mechanism of life, and is essential for growth and development. Bacteria and Eukarya have different mechanisms for cell division, which have been studied in detail. In contrast, cell division in Archaea is still understudied, and its regulation is poorly understood. Interestingly, different cell division machineries appear in the Archaea, with the Euryarchaeota using a cell division apparatus based on the tubulin-like cytoskeletal protein FtsZ, as in bacteria. Here we identify the small protein CdrS as essential for survival and a central regulator of cell division in the Euryarchaeon *Haloferax volcanii*. CdrS also appears to coordinate other cellular pathways including synthesis of signalling molecules and protein degradation. Our results show that CdrS plays a sophisticated role in cell division, including regulation of numerous associated genes. These findings are expected to initiate investigations into conditional regulation of division in archaea.

## Introduction

Cell division is a central aspect of the biology of all living organisms. In almost all bacteria, cell division is mediated by a ring-like division complex, or divisome, assembled around FtsZ, the ancestral homolog of eukaryotic tubulins that form the network of microtubules as part of the cytoskeleton (1). Bacterial FtsZ polymerizes into dynamic filaments and then assembles the contractile ‘Z-ring’ structure around the middle plane of the cell to constrict during cell division (2), which performs multiple functions, including recruiting divisome proteins to the division site (2), effecting membrane constriction (3), and guiding cell wall synthesis (4, 5). It is well known that bacteria can tightly coordinate division with growth rate to accurately duplicate their genomes and to homeostatically regulate their cell sizes (6, 7). A number of metabolic enzymes/pathways have been shown to directly regulate division in response to nutrient/metabolic status, by modulating the activity and assembly of FtsZ to ensure faithful cell division (8-10). Bacteria also regulate cell division in response to stresses including DNA damage. In *E. coli* the DNA damage or “SOS response” induces the expression of many genes, including FtsZ-specific inhibitors that block division. After the SOS response is turned off, the cell division inhibitor is degraded by proteases, allowing for cell division to resume (11).

Almost all bacterial species contain only one FtsZ (12), whereas many archaea carry two distinct FtsZ (FtsZ1 and FtsZ2) (13, 14). *Haloferax volcanii* has been proposed as a powerful model for understanding archaeal cell division and morphology (14-16). A recent study used *H. volcanii* to identify a new mechanism of FtsZ-based cell division; FtsZ1 has an initial scaffold-like function to stabilize the machinery controlling cell division and shape, and FtsZ2 is more actively involved in division constriction (16). However, how archaea regulate cell division in response to their environment and metabolism is not understood. A recent study has shown that metal micronutrients in the growth medium affect the cell size and shape of *H. volcanii*, suggesting a potential link between nutrient availability and the regulation of cell division (17). Another regulator of cell division in archaea may be a second messenger, <u>c</u>yclic <u>di</u>-<u>a</u>denylate <u>m</u>ono<u>p</u>hosphate (c-di-AMP), which was shown to be essential and tightly regulated in *H. volcanii* (18). Alteration of c-di-AMP levels in *H. volcanii* changed the average cell size in a low-salt medium, implying a function of c-di-AMP in regulation of cell size and division.

The current annotation of the *H. volcanii* genome shows 4,105 annotated protein-encoding genes (HaloLex 26.11.19) (19) and 316 of these open reading frames (ORFs) code for small proteins of 70 amino acids^1^ or less (20). Recent data show that proteins smaller than 70 amino acids are common and fulfil important functions in Bacteria and Eukarya (reviewed in (21-27)). Until more recently, sequences encoding such small proteins had long been overlooked and omitted from functional analyses (22, 28). Small proteins of Archaea have only been addressed in a few studies, which have implicated them in the regulation of nitrogen metabolism, protein degradation, oxidative stress response and sulphur metabolism (29-37). This limited body of information suggests a great potential of small proteins in the regulation of archaeal metabolism and biology. Quantitative proteome analysis of *H. volcanii* in standard as well as two stress conditions identified 60 of the annotated 316 small proteins predicted in *H. volcanii* (20). We have identified a small protein-encoding gene (HVO_0582), adjacent to *ftsZ2* in *H. volcanii*, which is highly transcribed according to a transcription start site analysis (38) (Figure 1). Based on the results reported here and in accordance with the concurrent study of its homologue from *Halobacterium salinarum* (39), we termed this protein CdrS (<u>c</u>ell <u>d</u>ivision <u>r</u>egulator - <u>s</u>hort).

**Figure 1.**
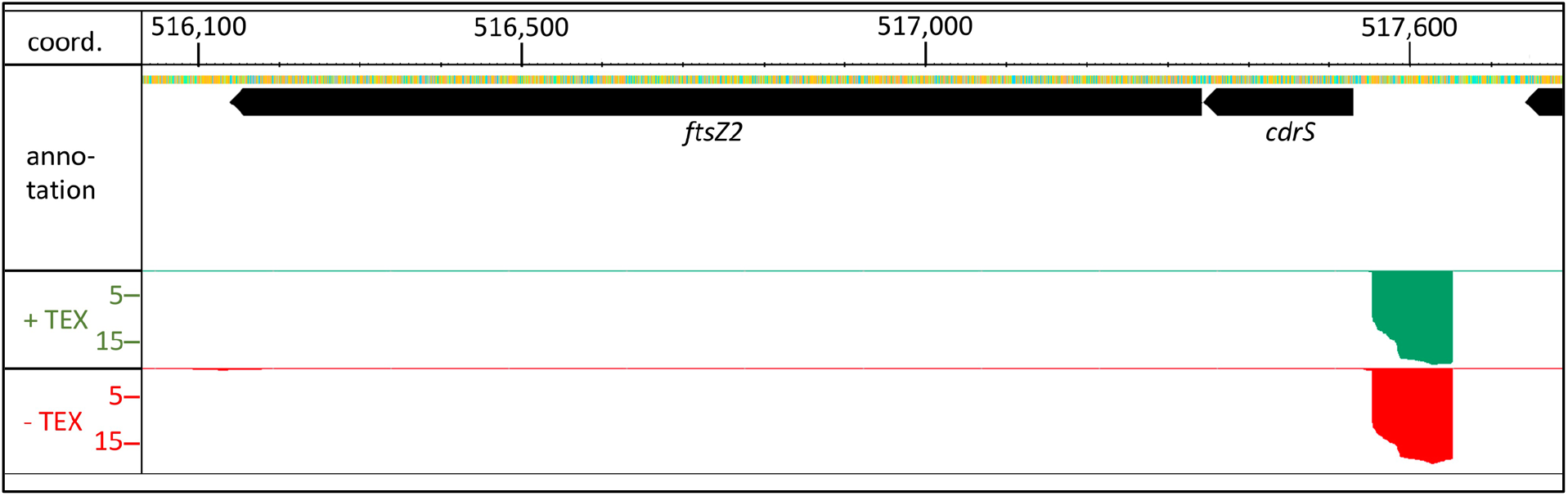
Genomic location of the *cdrS* gene. The gene for the small protein CdrS is encoded upstream of the *ftsZ2* gene encoding a homolog of the bacterial cell division protein FtsZ. The genes are separated by only two nucleotides. dRNA-Seq data (38) (lower panels) reveal a strong promoter upstream of the *cdrS* gene (data was visualized with the Integrated Genome Browser (40)). Red signals (panel -TEX) represent reads from an RNA fraction containing all cellular RNAs (primary transcripts as well as processed transcripts, read depth × 10^3^). Green signals (panel +TEX) represent reads from an RNA sample treated with terminator 5’ phosphate dependent exonuclease (TEX) (read depth × 10^3^), resulting in enrichment of primary transcripts. Comparison of both data sets allows determination of transcription start sites. The genome coordinates and the annotation are shown at the top.

The *cdrS-ftsZ2* locus is well conserved across Euryarchaeota, especially within the class of Halobacteria. Using *H. volcanii* as the model organism, we found that the *cdrS* gene is essential, and unlike *ftsZ2*, could not be deleted. As *cdrS* encodes a predicted transcription regulator, we used an integrative approach to investigate its functions by combining gene repression by CRISPR interference (CRISPRi), ChIP-Seq, transcriptomics, quantitative proteomics, and microscopy. Our data suggest that CdrS is a global transcriptional regulator, controlling *ftsZ* expression and genes linked to other metabolic and regulatory processes. This may allow cells to properly coordinate growth, division and metabolic activity.

## Results

### Repression of cdrS-ftsZ2 expression causes cell growth defects in H. volcanii

CdrS is predicted as a small protein of 61 amino acids with an isoelectric point of 9.5, which is very basic for a halophilic protein and might indicate that it binds to negatively charged molecules like nucleic acids. The *cdrS* open reading frame (ORF) is located three nucleotides upstream of the *ftsZ2* ORF suggesting that they might be transcribed together (Figure 1). The currently predicted function of CdrS is a transcriptional regulator containing the CopG/Arc/MetJ DNA-binding domain with a ribbon-helix-helix (RHH) motif. To our knowledge, similar small transcription factors have so far been only described for bacteria (41, 42).

To help uncover the biological functions of CdrS, we first attempted to generate a *cdrS* deletion mutant to observe its functional consequences. Several attempts to generate such a strain using the standard pop-in/pop-out method (43) proved unsuccessful, suggesting that *cdrS* is essential. Therefore, we employed CRISPRi to repress expression of *cdrS* (Supplementary Figure 1). The CRISPRi approach takes advantage of the endogenous CRISPR-Cas system of *H. volcanii* that can be harnessed to repress transcription. It was already successfully used in *H. volcanii* to repress expression of several genes (44, 45). Three different crRNAs were designed that bind to the promoter and transcription start site regions of *cdrS* and guide the endogenous Cascade complex to these sequences thereby preventing transcription initiation (Figure 2A). *H. volcanii* cells were transformed with plasmids for expression of the three crRNAs and northern blot analyses showed that in wild-type cells^2^ the bicistronic *cdrS*-*ftsZ2* mRNA was the predominant transcript (∼1,500 nt), and that crRNAs anti#1, anti#2 and anti#3 repressed its expression moderately, to an average of 93%, 76% and 60% of the wild-type, respectively (Figure 2B, Supplementary Figure 2A). The monocistronic *cdrS* mRNA (∼250 nt) was repressed by all three crRNAs (57%, 42% and 40% of the wild-type, respectively) (Figure 2B, Supplementary Figure 2B). Additional RNAs were also observed, which could be generated by cleavage of the longer bicistronic mRNA or by premature termination of transcription. All three CRISPRi strains showed reduced growth rates in comparison to the control strain expressing no crRNA (Figure 2C), with doubling times of 3.9±0.15 h (wild-type control, HV35 × pTA232), 4.2±0.07 h (*cdrS* CRISPRi#1), 8.4±0.4 h (*cdrS* CRISPRi#2) and 8.9±0.84 h (*cdrS* CRISPRi#3).

**Figure 2.**
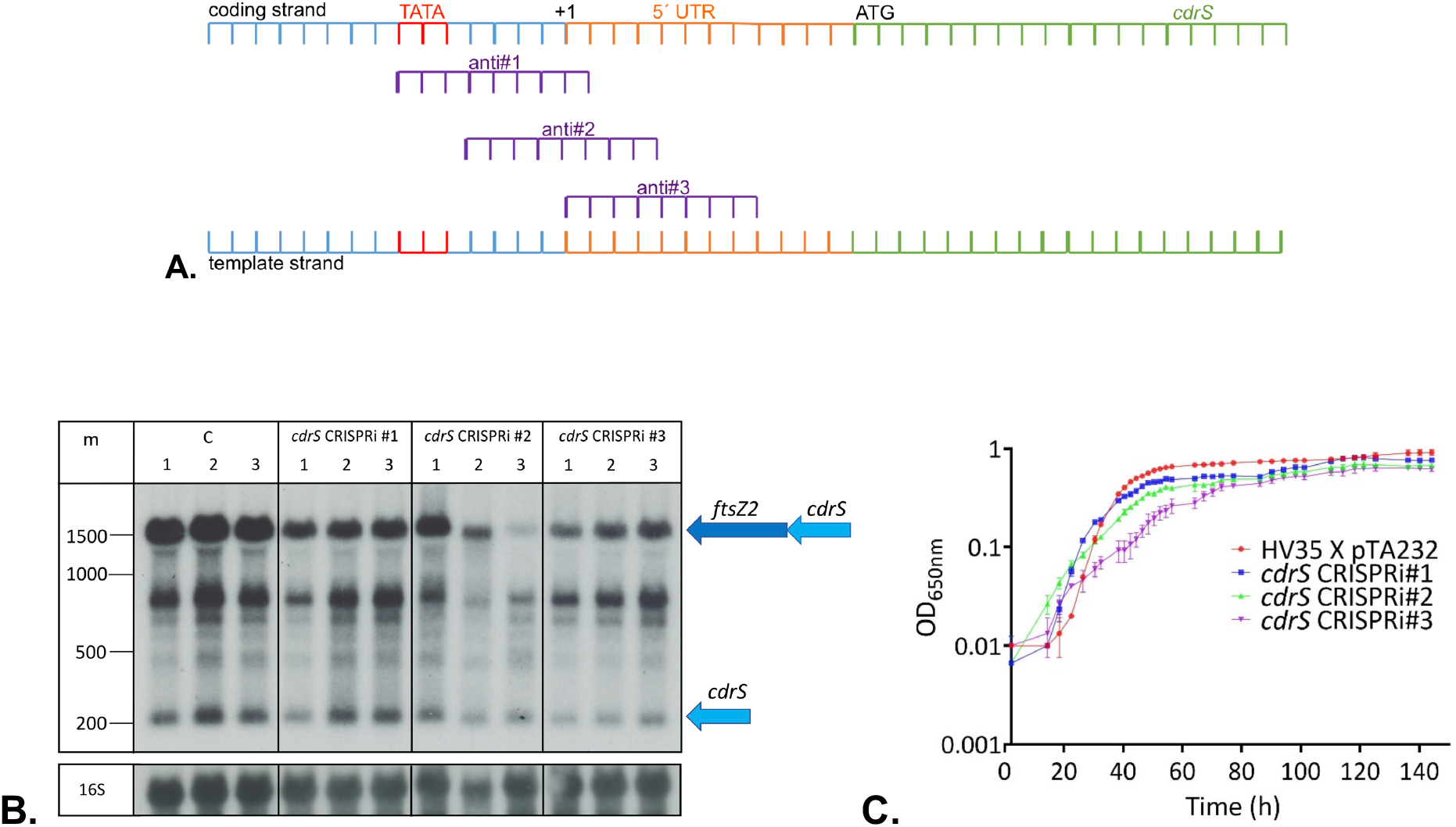
Repression of *cdrS-ftsZ2* and its effect on growth. A. Location of crRNAs directed against the *cdrS* gene. Both strands of the *cdrS* upstream region are shown. Three different crRNAs (anti#1-3) were designed to target the template strand in the promoter region and close to the transcription start site. The TATA box is shown in red, the transcription start site is indicated as “+1”, the 5’
s UTR region is shown in orange, the open reading frame in green and the ATG start codon is indicated. **B. Both *cdrS and ftsZ2 are repressed* by CRISPRi**. Hybridisation with a probe against the *cdrS* mRNA (upper panel) revealed the monocistronic *cdrS* mRNA (signal at about 250 nucleotides) as well as the bicistronic *cdrS*-*ftsZ2* mRNA (signal at about 1,500 nucleotides) and intermediate RNAs that may be degradation products. Lanes C: wild-type RNA (HV30 x pTA232): HV30 strain with pTA232 plasmid without insert; lanes *cdrS* CRISPRi #1, #2 and #3: HV30 strain expressing crRNAs anti#1, anti#2 and anti#3 (HV30 x pTA232-tele-0582anti#1-3). Experiments were done in biological triplicate (lanes 1, 2, 3 in each case). The lower panel was hybridised with a probe against the 16S rRNA. An RNA size marker is given at the left in nucleotides (lane m). The mono- and the bicistronic mRNA are shown at the right schematically. **C. Growth of the *cdrS* CRISPRi strains is impaired**. Growth of wild-type *Haloferax* strain (HV35 x pTA232) was compared to growth of the CRISPRi strains expressing crRNA anti#1, crRNA anti#2 or crRNA anti#3 (*cdrS* CRISPR#1-#3). Analyses were done in triplicate; standard deviations are shown as bars. X-axis shows the time of growth (in hours), y-axis shows the OD semi-logarithmically.

### Repression of cdrS-ftsZ2 causes changes in the transcriptome

To determine whether *cdrS-ftsZ2* repression influences the expression of other genes we compared transcriptomes of the CRISPRi strain *cdrS* CRISPRi#2 and the wild-type strain (Supplementary Table 1). The transcriptome data confirmed repression of *cdrS* (log_2_ -2.5) and *ftsZ2* (log_2_ -2.6). In addition, they revealed expression differences for 97 genes (with a log_2_ fold change lower and higher than -0.7 and +0.7, respectively).

Thirty-five genes are upregulated (Supplementary Table 1A) and 62 genes downregulated (Supplementary Table 1B). Thirty genes encoding secreted, membrane and cell surface proteins were differentially expressed (12 upregulated and 18 downregulated), suggesting significant changes to the cell envelope. Other upregulated genes included one involved in transport, four in transcription regulation (which could mediate some effects of *cdrS* repression), two in signal transduction, two in DNA maintenance and repair, three in metabolism, one in branched side-chain amino acid biosynthesis, and 11 with unknown functions (Supplementary Table 1A).

Four downregulated genes are implicated in the cell cycle and division, including *ftsZ1, ftsZ2, sepF*, and a *smc* (structural maintenance of chromosomes) homologue. Other downregulated genes are implicated in transport of branched-chain amino acids and sugars, transcription, cobalamin (vitamin B12) biosynthesis, amino acid metabolism (arginine/lysine) and general metabolism (Supplementary Table 1B).

Since the CRISPRi approach represses both *cdrS* and *ftsZ2*, we next aimed to identify genes that are specifically regulated upon *cdrS* repression only, by complementing the CRISPRi strain with either the *ftsZ2* gene alone or the *cdrS-ftsZ2* genes together. For northern blot analyses we selected two genes which were found to be regulated in the *cdrS* CRISPRi transcriptome: the upregulated genes HVO_B0192-HVO_B0193 and the downregulated gene HVO_0739. Northern blots with RNAs from wild-type (HV30 x pTA232 x pTA409), *cdrS* CRISPRi (HV30 x pTA232-tele-0582anti#2 x pTA409) and complemented strains (HV30 x pTA232-tele-anti#2 x pTA409*ftsZ2*, HV30 x pTA232-tele-anti#2 x pTA409*ftsZ2-cdrS*) were hybridised with probes against the selected mRNAs (Figure 3). Consistent with the transcriptome results the genes HVO_B0192-HVO_B0193 were found to be upregulated in the CRISPRi#2 strain complemented with *ftsZ2* only (Figure 3). Likewise, HVO_0739 was confirmed to be downregulated in the CRISPRi cells complemented with only *ftsZ2*.

**Figure 3.**
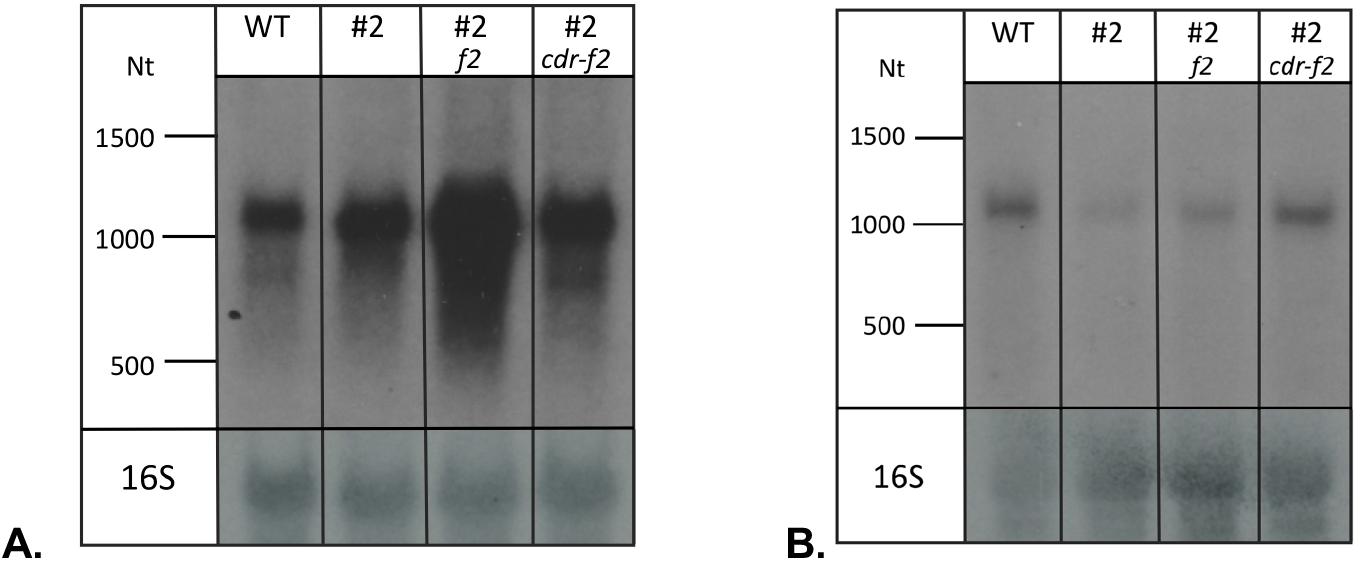
Northern analyses reveal specific effects of repression of *cdrS* only. RNA was analysed from wild-type cells (lane WT) (HV30 x pTA232 x pTA409), *cdrS* CRISPRi#2 cells (lane #2) (HV30 x pTA232-tele-0582anti#2 x pTA409) and *cdrS* CRISPRi#2 cells complemented with *ftsZ2* (lane #2 *f2*)(HV30 x pTA232-tele-0582anti#2 x pTA409-pfdx-HVO_0581-nat.t (*ftsZ2*)) and *cdrS*-*ftsZ2* (lane *cdr-f2*) (HV30 x pTA232-tele-0582anti#2 x pTA409-pfdx-HVO_0582-HVO_0581-nat.t (*cdrS*-*ftsZ2*)). **A**. Hybridisation with a probe against HVO_B0192 and HVO_B0193, two genes that are upregulated in CRISPRi cells, confirmed upregulation (lane #2) and upregulation is even more prominent in CRISPRi cells complemented with *ftsZ2 only*. Both genes are transcribed together into an approximately 1,077 nucleotides mRNA. **B**. Hybridisation with a probe against HVO_0739, a gene that is downregulated in the CRISPRi strain, confirmed that the mRNA is downregulated in CRISPRi cells (lane #2). The mRNA is also downregulated in CRISPRi cells complemented with *ftsz2* (lane *f2*). Complementation with both genes results in wild-type mRNA levels (lane *cdr-f2*). The gene is transcribed into a monocistronic mRNA of about 1,052 nucleotides. Both membranes have also been hybridised with a probe against the 16S rRNA (lower panel).

Taken together, the data indicated that upregulation of HVO_B0192-HVOB0193 and downregulation HVO_0739 were due specifically to *cdrS* repression.

### Repression of cdrS-ftsZ2 with CRISPRi causes multiple changes to the proteome

We next compared the soluble and insoluble fractions of the wild-type and CRISPRi (*cdrS* CRISPRi#2) strains by quantitative proteomics. Previous proteome analyses of *H. volcanii* have shown that standard mass spectrometry techniques are biased against the detection of small proteins (20, 47). However, the CdrS protein was identified in two of the three wild-type replicates (with one peptide each), but with detection in only two replicates it did not meet our criteria for listing in Supplementary Table 2. CdrS was not detected in any of the three CRISPRi strain replicates, consistent with repression of the *cdrS* gene. Thirty-four proteins were found to be more abundant or only found in CRISPRi cells, including two hypothetical transmembrane proteins, two involved in transport, four in translation, five in carbohydrate metabolism, one in central carbon metabolism (acetyl-CoA synthetase), one in arginine biosynthesis, one in lipid metabolism (isoprenyl diphosphate synthase), three in DNA replication and repair, eleven in general metabolism, and four of unknown functions (Supplementary Table 2). Conversely, 23 proteins were found to have lower abundance or were absent from the CRISPRi cells, including six transmembrane proteins, two involved in transport, one 30S ribosomal protein S15, two isocitrate lyase regulator-type (IclR-type) transcriptional regulators, one in carbohydrate metabolism, two in amino acid metabolism, four in general metabolism, one photolyase homologue (*phr1*) for DNA repair, two in cell division (FtsZ2 and SepF), and two of unknown functions (Supplementary Table 2). The change in abundance of proteins implicated particularly in metabolism, metabolite transport and regulation, and cell division, suggests that CdrS might be involved in coordinating aspects of metabolism and growth with division.

### CdrS associates with a specific DNA motif in vivo

Since the predicted function for CdrS is a transcriptional regulator, we identified DNA binding sites of CdrS *in vivo* using chromatin-immuno-precipitation and DNA sequencing (ChIP-Seq). Eighteen binding sites were revealed by ChIP-Seq (Table 1 and Figure 4A.), which allowed the identification of a specific binding site motif (Figure 4C.). For 15 of the 18 sites the motif is located between 110 to 41 nucleotides upstream of the transcription start site (TSS) of a gene (Table 1). One binding site is located upstream of two closely spaced TSS (−57 and -74 nucleotides upstream), two binding sites are overlapping with a TSS, and for one binding site no TSS is present under the standard growth conditions. Two binding sites were found in close proximity on opposite strands of two divergent genes, *hisB* (HVO_2986) and HVO_2987 (Supplementary Figure 3). CdrS binding sites were only detected on the main chromosome and not on the mini-chromosomes pHV1, pHV3 and pHV4.

**Table 1.**
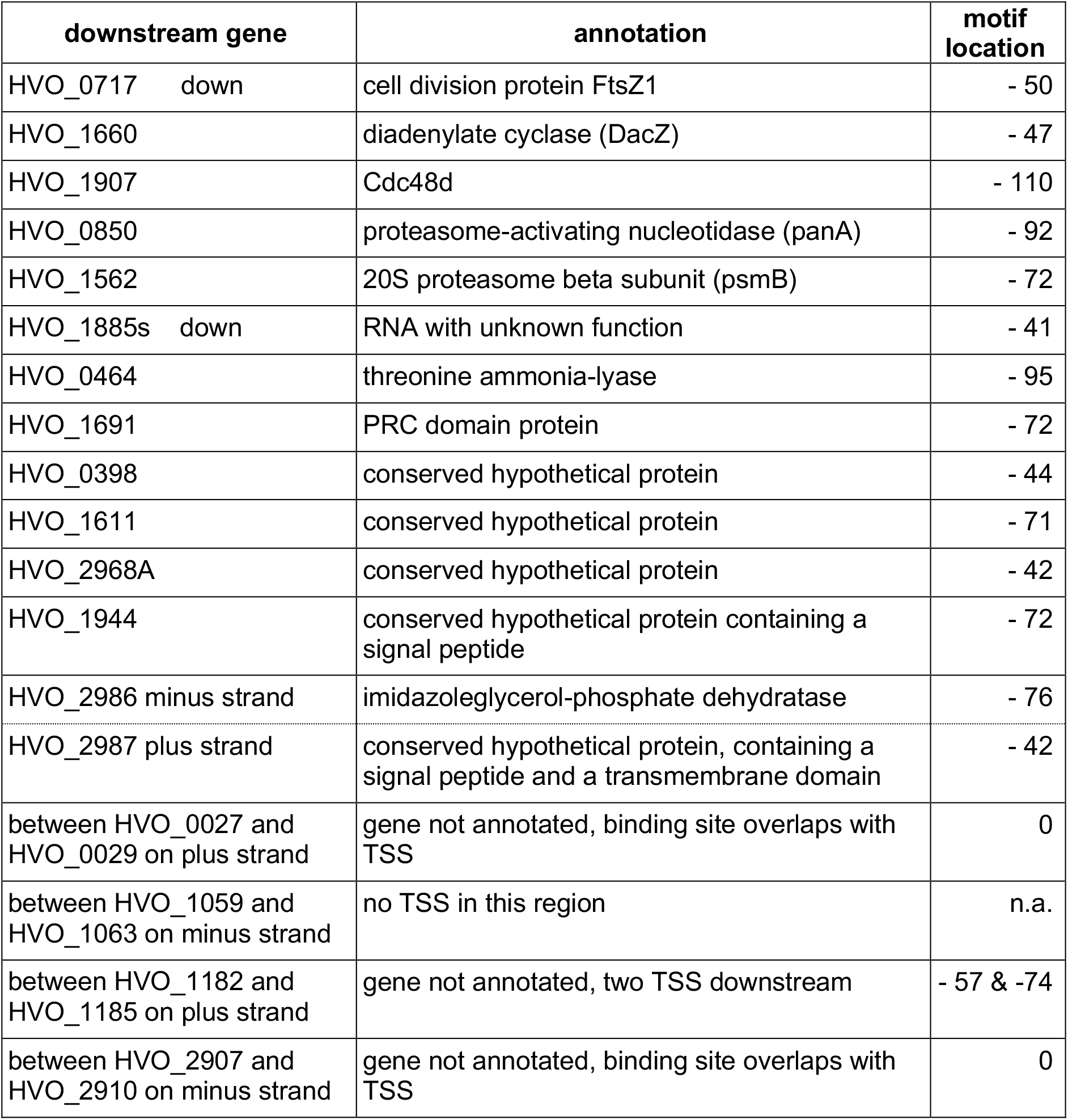
DNA binding sites identified for CdrS. ChIP-Seq showed that CdrS binds to 18 different sites in the chromosome. For one location the binding site is present on both strands (HVO_2986 and HVO_2987). Column “downstream gene” lists the gene located downstream of the target site and whether the gene is downregulated in the CRISPRi strain (indicated by “down”)(Supplementary Table 1B), column “annotation” shows the annotation for that gene, and column “motif location” shows the distance of the identified binding motif (counting from the 10^th^ nucleotide of the motif) to the transcription start site of each gene.

**Figure 4.**
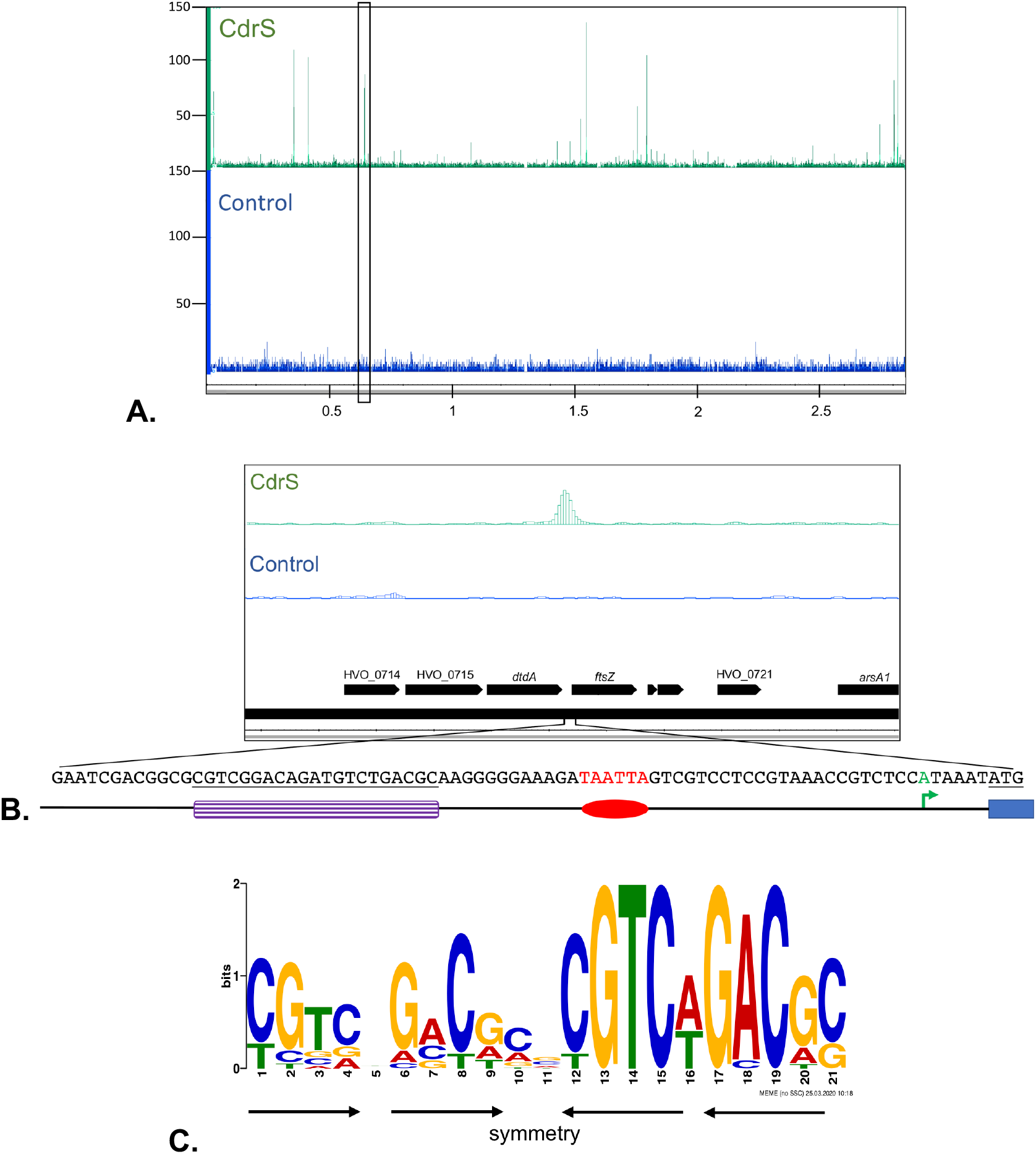
Identification of the CdrS DNA binding motif. ChIP-Seq was employed to identify binding sites of CdrS. Panel “control”: ChIP-Seq analysis with the control sample (blue); “CdrS”: ChIP-Seq with the CdrS protein (green). **A. CdrS binds to eighteen locations in the genome**. The ChIP-Seq data are shown for the complete main chromosome (2.8 Mb). The chromosomal region highlighted by the black rectangle is shown enlarged in panel B. Read numbers are shown at the left; the chromosome coordinates are shown in Mb at the bottom. **B. CdrS binding site upstream of *ftsZ1***. The chromosomal region highlighted by the black rectangle in panel A is located upstream of the *ftsZ1* gene. Annotated genes are shown at the bottom, the sequence upstream of the *ftsZ1* gene is enlarged below the genome annotation. The CdrS binding motif is shown in purple, the promoter is shown in red, the TSS is shown in green and the start of the gene (ATG) is underlined and shown as blue box. **C**. The conserved DNA binding motif identified using MEME-ChIP (48) shows notable symmetry (arrows), which might reflect the binding of CdrS multimer.

CdrS bound upstream of the *ftsZ1* gene (Figure 4B.), implicating CdrS in the regulation of a gene involved in cell division (16). According to the transcriptome data, the *ftsZ1* gene is moderately repressed in CRISPRi cells (HVO_0717, log_2_ fold change -1.3) (Supplementary Table 1B), suggesting that CdrS may activate *ftsZ1* transcription in wild-type cells. Proteome data of the CRISPRi strain showed an FtsZ1 abundance that appeared slightly lower than in the wild-type (log_2_: -1.2) but with a p-value of 0.13 it did not pass the parameter threshold. However, western blot analyses with antibodies raised specifically against FtsZ1 and FtsZ2 were consistent with the above data; FtsZ2 was not detected in CRISPRi cells and FtsZ1 concentrations were reduced (Figure 5F.).

**Figure 5.**
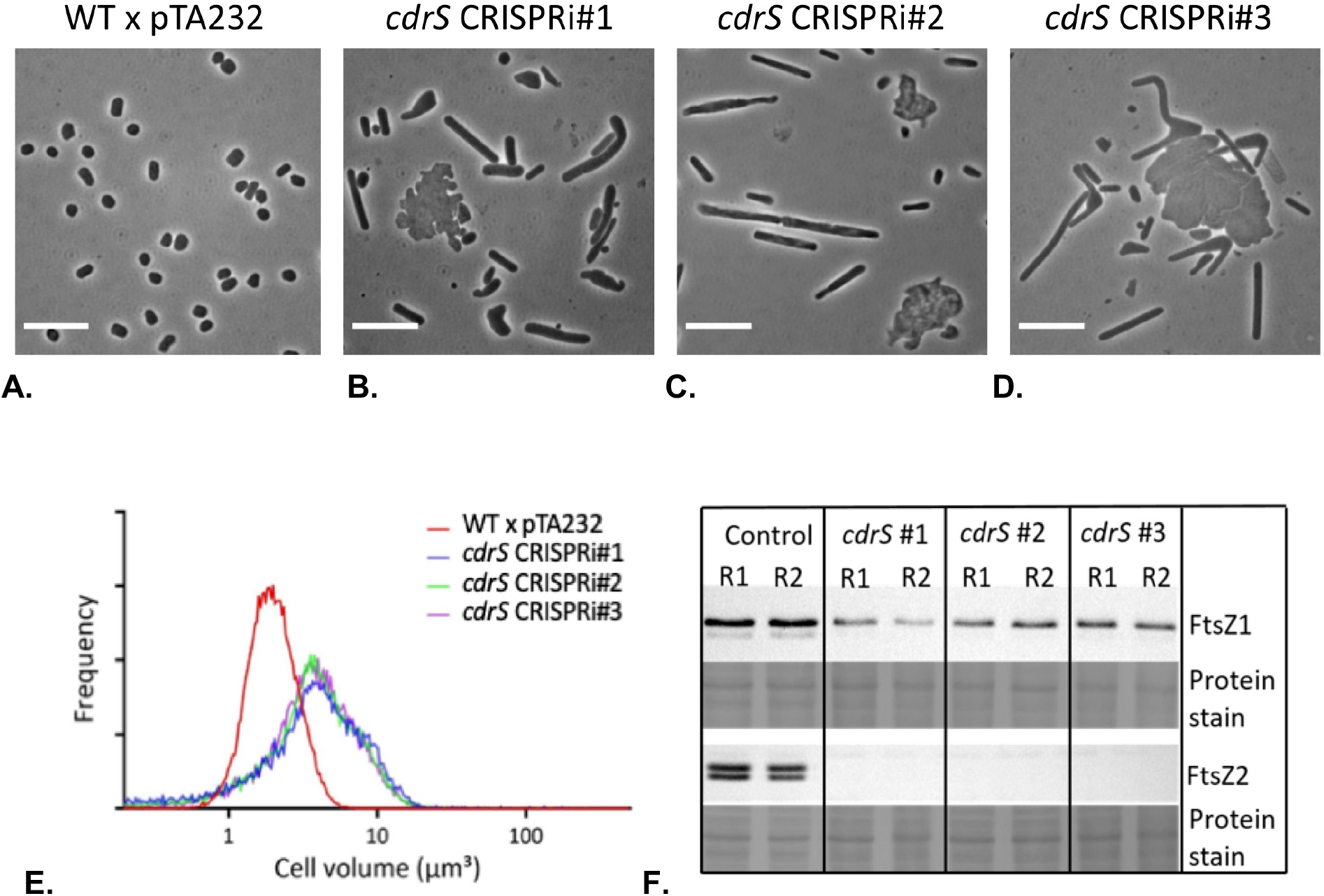
CRISPRi repression of *cdrS-ftsZ2* results in defects in cell division. **A**.**-D**. Phase-contrast images of **A**. the wild-type strain without crRNA expression: WT x pTA232 (HV35 x pTA232), and three *cdrS* repression strains expressing the three different crRNAs: **B**. *cdrS* CRISPRi#1 (HV35 x pTA232-0582anti#1); **C**. *cdrS* CRISPRi#2 (HV35 x pTA232-0582anti#2); and **D**. *cdrS* CRISPRi#3 (HV35 x pTA232-0582anti#3). All strains were sampled during steady mid-log phase in Hv-MinTE medium supplemented with 50 µg/ml uracil and 0.04 mM Trp. Scale bars, 10 µm. **E**. Coulter cytometry cell volume distributions obtained from the samples shown in panel A-D. Cell volume is shown at the x-axis and frequency (relative fraction of total cells) on the y-axis. **F**. Western blot analysis of FtsZ1 and FtsZ2 expression levels for control (HV35 x pTA232) and three *cdrS* repression strains (*cdrS* #1: HV35 x pTA232-0582anti#1; *cdrS* #2: HV35 x pTA232-0582anti#2; *cdrS* #3: HV35 x pTA232-0582anti#3). Total protein pre-staining of each membrane (with Ponceau S) is shown as a loading control. R1 and R2: two independent biological replicate protein samples. Data displayed are representative of at least two technical replicate experiments.

Another gene targeted by CdrS was the gene for diadenylate cyclase, which is essential and generates the signalling molecule c-di-AMP in *H. volcanii* (18). CdrS also binds upstream of the promoter for an RNA gene of unknown function, HVO_1885s. HVO_1885s was downregulated in CRISPRi cells according to the transcriptome data (log_2_: -2.2, Supplementary Table 1B) and therefore might normally be activated by CdrS. Further target genes encode proteins that are involved in proteasome activity (Cdc48d (49), PanA (50, 51) and PsmB (50, 51)). In addition, CdrS binds to seven genes encoding proteins and an RNA with unknown functions, as well as to three regions upstream of unannotated potential genes.

### Repression of cdrS-ftsZ2 with CRISPRi has a severe effect on cell size and morphology

Microscopic analyses revealed that the three CRISPRi strains showed substantial changes in cell size and morphology featuring giant and misshapen plate-like cells as well as long filamentous cells during mid-logarithmic phase (Figure 5B-D and Supplementary Figure 4). The giant cells are a hallmark of a cell division defect, since cells grow but division is delayed or fails, resulting in overgrowth. As *H. volcanii* can exist as rods or plate-shaped cells in culture, the filamentous and giant-plate cell types in the CRISPRi strains are expected to be the result of a cell division defect in these two cell morphotypes, respectively (16). The three CRISPRi strains had very similar cell-size defects when compared to the wild-type cells (Figure 5A-E.) which were consistent throughout the growth phases (Supplementary Figure 5) and showed similar cell morphology profiles (Supplementary Figure 4 and Supplementary Table 3). Western blotting showed that repression of *cdrS* decreased FtsZ1 concentrations slightly and FtsZ2 levels drastically (Figure 5F). The effect on FtsZ1 varied somewhat for the three CRISPRi strains (in *cdrS* CRISPRi#1 FtsZ1 was ∼35% decreased compared to wild-type; *cdrS* CRISPRi#2 ∼69%; *cdrS* CRISPRi#3 ∼76%), whereas FtsZ2 was strongly depleted in all cases (Figure 5F). Together, these data implicate CdrS in the regulation of cell division, and that it directly or indirectly influences cell shape.

### Cell division defects are rescued by complementation with both cdrS and ftsZ2

To determine whether the cellular effects we observed in Figure 5 were due to repression of *cdrS* and *ftsZ2* together, or either one, we complemented the CRISPRi strain (*cdrS* CRISPRi#3) with plasmids expressing the *cdrS* gene alone, the *ftsZ2* gene alone, or both genes together. Only complementation with both genes together rescued the cell morphology and division defects (Figure 6 and Supplementary Figure 6). Western blotting showed that the complementation with *cdrS* alone did not restore FtsZ2 levels and the FtsZ2 level was restored when *ftsZ2* was included on the complementation plasmid, as expected (Figure 6G). These data suggest that CdrS is important for cell division, independently of FtsZ2.

**Figure 6.**
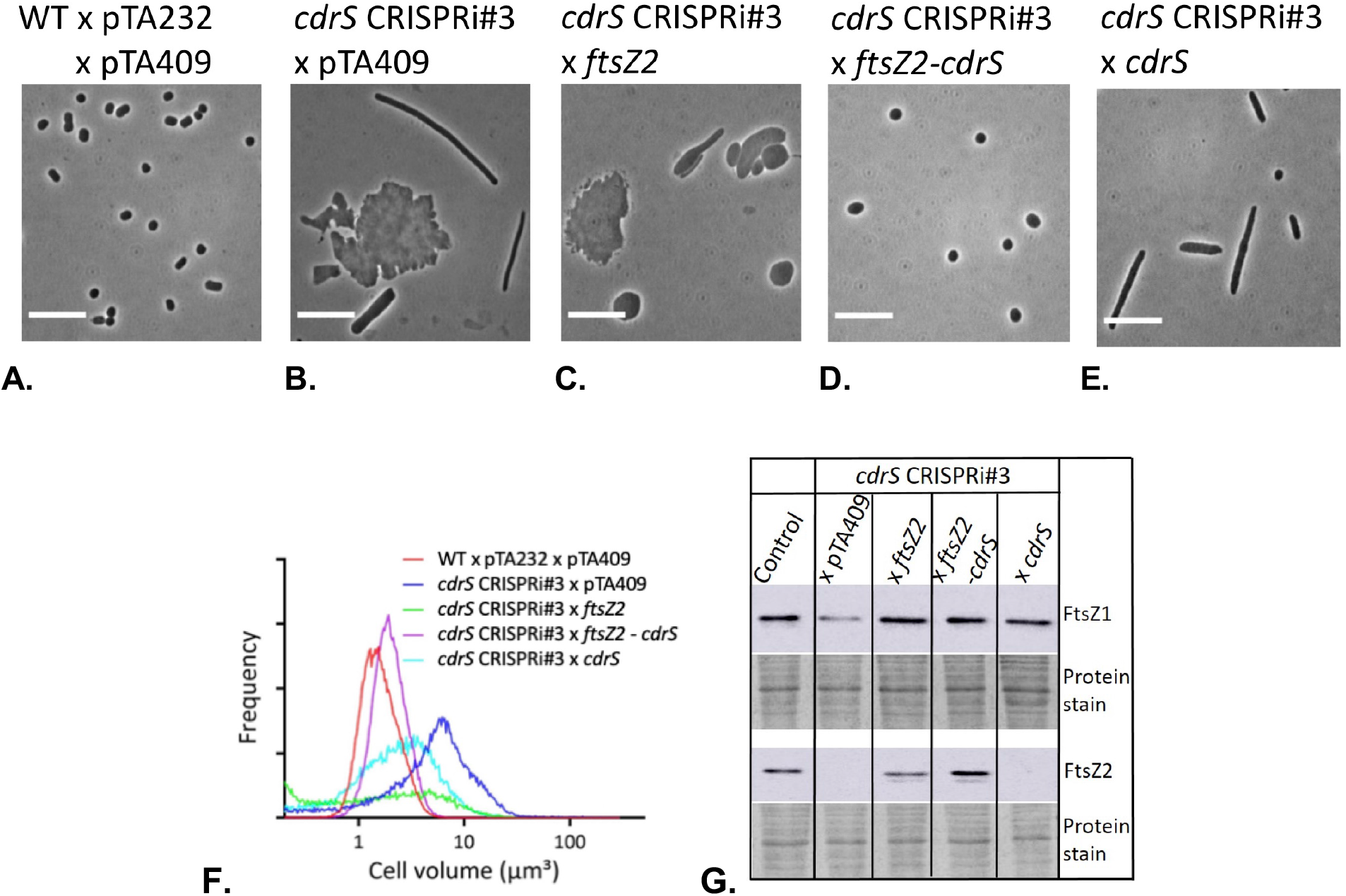
Complementation of CRISPRi strains. **A**.**-E. Phase contrast micrographs**. Cell size in the CRISPRi#3 strain appeared normal with supplemental expression of both *ftsZ2* and *cdrS* but not during expression of *ftsZ2* or *cdrS* individually. Phase-contrast images of **A**. wild-type vector-only as the control for complementation (HV35 x pTA232 x pTA409), **B**. *cdrS* CRISPRi#3 vector only (HV35 x pTA232-0582anti#3 x pTA409), **C**. *ftsZ2* complementation of CRISPRi#3 cells (HV35 x pTA232-0582anti#3 x pTA409_*ftsZ2*), **D**. *ftsZ2*-*cdrS* double complementation of CRISPRi#3 cells (HV35 x pTA232-0582anti#3 x pTA409*ftsZ2-cdrS*), and **E**. *cdrS* complementation of CRISPRi#3 cells (HV35 x pTA232-0582anti#3 x pTA409*cdrS*). Scale bars, 10 µm. **F. Coulter cell volume analysis**. Coulter cell volume analysis of the complementation effects of *cdrS* CRISPRi#3, samples as listed in A.-E. **G**. Western blot analysis of FtsZ1 and FtsZ2 expression levels for control (HV35 × pTA232 × pTA409), *cdrS* CRISPRi#3 vector only, *ftsZ2* complementation of *cdrS* CRISPRi#3, *ftsZ2-cdrS* double complementation of *cdrS* CRISPRi#3, and *cdrS* complementation of *cdrS* CRISPRi#3. Total protein pre-staining of each membrane (with Ponceau S) is shown as a loading control. Data displayed are representative of two independent experiments.

### Overexpression of cdrS induces cell size and morphological defects

We next sought to identify any effects of overexpression of *cdrS, ftsZ2*, or both genes together, from a constitutive strong promoter in a wild-type background. Interestingly, overexpression of *cdrS* alone had a clear effect on cell size and morphology, showing some elongated (2.3 %), large and misshapen cells (45 %), as well as some wild-type like cells (40.5 %) (Figure 7D and E, Supplementary Figure 7 and 8, Supplementary Table 3), consistent with mis-regulated division. In contrast, FtsZ2-only overexpression showed slightly smaller cells than wild-type cells (Figure 7B and E, Supplementary Figure 7 and 8), consistent with results using an inducible promoter (16), termed hyper-division. Furthermore, overexpression of both *cdrS* and *ftsZ2* together resulted in wild-type like cells that had significantly smaller cell size than both the wild-type and *ftsZ2*-only overexpression strains (Figure 7C and E, Supplementary Figure 7 and 8).

**Figure 7.**
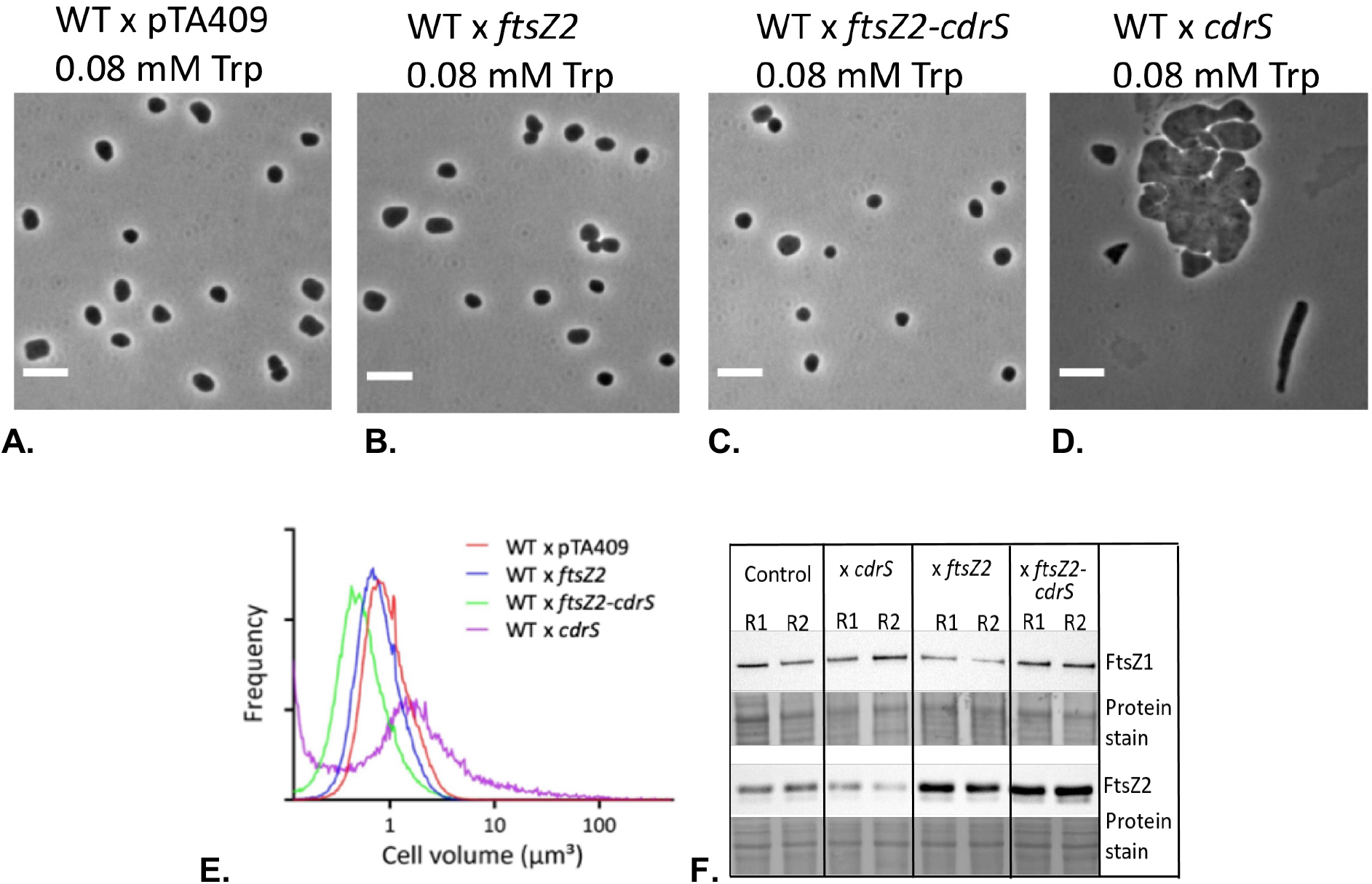
Overproduction of FtsZ2 and/or CdrS in wild-type background differentially affected cell division. **A**.**-D.** Phase contrast micrographs. **A**. Wild-type (HV35 x pTA409), **B**. FtsZ2 overexpression (HV35 x pTA409*ftsZ2*), **C**. FtsZ2-CdrS double overexpression (HV35 x pTA409*ftsZ2-cdrS*), and **D**. CdrS overexpression (HV35 x pTA409*cdrS*). Compared to wild-type, both FtsZ2 single- and FtsZ2-CdrS double overexpression produced cells having wild-type morphology, whereas CdrS overexpression resulted in elongated/enlarged cells. Scale bars, 5 µm. **E**. Coulter cell volume analysis. Cells overexpressing FtsZ2 produced cells with slightly smaller size than wild-type, and FtsZ2-CdrS double overexpression produced significantly smaller cells reflecting hyper-cell division, while CdrS overexpression resulted in significantly larger cells indicating inefficient or mis-regulated cell division. **F**. Western blot analysis of FtsZ1 and FtsZ2 expression levels for control (HV35 x pTA409), CdrS overexpression, FtsZ2 overexpression, and FtsZ2-CdrS double overexpression. Total protein pre-staining of each membrane (with Ponceau S) is shown as a loading control. R1 and R2: two independent biological replicate protein samples. Data displayed are representative of at least two technical replicate experiments.

Western blot analysis of samples taken during *cdrS* overexpression indicated very little difference in the level of FtsZ2 (∼0.7 fold) compared to wild-type control, whereas the FtsZ1 level increased ∼1.7 fold (Figure 7F), consistent with the CRISPRi results that suggest CdrS moderately promotes *ftsZ1* expression (Figure 5). During *ftsZ2* overexpression, FtsZ1 levels were similar to wild-type, whereas FtsZ2 levels were as expected substantially higher (∼2.3 fold) than wild-type. FtsZ2 levels increased to ∼2.9-fold higher than the wild-type in the *cdrS*-*ftsZ2* double overexpression strain (Figure 7F), which might account for the additional stimulation of division observed in this strain (Figure 7E). Finally, FtsZ1 increased ∼1.5-fold vs during double *cdrS*-*ftsZ2* overexpression, consistent with *cdrS* overexpression and CRISPRi repression results.

## Discussion

Microscopic analyses of *H. volcanii* strains undergoing CRISPRi-mediated repression of the small gene, *cdrS* (HVO_0582), supported by results obtained with complemented strains, showed that repression of *cdrS* expression alone has a severe impact on cell size and morphology; the most obvious defect at the cellular level appears to be in the regulation or mechanism of cell division. This is supported by the finding that the *cdrS* gene is co-transcribed with a cell division gene, *ftsZ2*. ChIP-Seq revealed 18 CdrS binding sites, including one upstream of *ftsZ1*, another homolog involved in division (15). During CRISPRi-mediated repression of *cdrS, ftsZ1* was moderately downregulated, and strong down regulation was seen for other genes that might be involved in cell division including HVO_0392 (encoding a homolog of the bacterial SepF division protein), and HVO_0739 (predicted membrane protein) (Table 2). Archaeal SepF was recently identified as an anchor for the division ring at the cytoplasmic membrane (52, 53). The identified differential abundance of other proteins/genes when *cdrS* is repressed, involved in cell surface/membrane, transport, lipid metabolism and carbohydrate metabolism (possible glycosylation), suggests that CdrS may be a global regulator that controls division and other envelope-related functions in response to nutritional or metabolic changes.

**Table 2.** Genes and proteins affected in both the transcriptome and proteome of CRISPRi cells. Three genes were found downregulated in the transcriptome and their protein products were found with lower abundance or “off” in the proteome.

| gene | protein | fold change log<sub>2</sub> |  |
| --- | --- | --- | --- |
|  |  | Transcriptome | Proteome |
| HVO_0581 | FtsZ2 | -2.6 | -3.2 |
| HVO_0392 | probable SepF protein | -2.2 | -3.3 |
| HVO_0739 | hypothetical protein | -3.3 | off |

Our results strongly suggest that CdrS functions as a transcription regulator. The majority of archaeal transcription factors have a helix-turn-helix (HTH) motif, only a few contain the RHH domain (54). To our knowledge similar small transcription factors have so far been only described for bacteria (41, 42). The binding location is usually an indicator how a transcription factor acts: those activating transcription typically bind upstream of promoters (54), whereas binding at or downstream of the promoter usually inhibit transcription by preventing recruitment of the RNA polymerase (54). CdrS binds upstream of the promoters of *ftsZ1* and HVO_1885s which are downregulated in the CdrS depletion strain, thus consistent with CdrS normally acting as transcriptional activator for these genes. CdrS also binds upstream of the *dacZ* gene that encodes diadenylate cyclase (DacZ), which synthesizes the second messenger molecule c-di-AMP. The *dacZ* gene is essential in *H. volcanii*, and overexpression of *dacZ* was lethal, indicating its central importance in cells, and the need for tight regulation (18). The targets in *H. volcanii* for c-di-AMP signaling are currently unknown; we speculate that c-di-AMP regulation via CdrS might play a role in coordinating metabolic processes with the cell division or other envelope-related mechanisms. Consistent with this, our findings implicate CdrS as an activator of vitamin B12 (cobalamin) biosynthesis in *H. volcanii* (Supplementary Table 1B), which has previously been noted to be under the regulatory network of the transcriptional regulator TrmB, a regulator of sugar metabolism in *H. salinarum* (55). Expression of the CdrS homolog in *H. salinarum* is itself regulated in response to oxidative stress (56). These results suggest that CdrS could take part in regulating division and envelope-related functions in response to multiple global response pathways.

We also found CdrS binding sites upstream of three genes related to proteasome function (*cdc48d, panA* and *psmB*). The gene *psmB* encodes the β subunit of the 20S proteasome in *H. volcanii*, which consists of proteins α1, β, and α2. The two proteasome-activating nucleotidases (PanA and PanB) are closely related to the regulatory particle AAA-ATPases (Rpt) of eukaryotic 26S proteasomes (50, 51). Cdc48-like proteins appear universal among archaea and are linked to the function of the 20S proteasome in archaea (57). It has recently been shown in *Sulfolobus* that the activity of the proteaosome is required for cells to divide and initiate the next round of DNA replication, thereby revealing a connection between the proteaosome and cell division (58). It is possible that CdrS regulates a similar connection between them in *H. volcanii*. As the FtsZ-based cell division apparatus is dispensable in *H. volcanii* (15), the essential function of the *Haloferax* CdrS might lie in regulation of the essential DacZ and proteasome proteins. Interestingly, the CdrS homolog in *H. salinarum* (HbaCdrS, VNG0194H), which is also encoded in an operon together with FtsZ2, can be deleted (56); *H. salinarum* CdrS appears not to be involved in regulation of DacZ and proteasome proteins, which might explain its non-essentiality.

We noted a general low correlation between ChIP-Seq, proteome and transcriptome datasets. A high correlation between the ChIP-Seq data and the Omics data is not to be expected, since ChIP-Seq data reveal the binding activity of only CdrS whereas transcriptome and proteome results are due to repression of both CdrS and FtsZ2. Low correlation between transcriptome and proteome data has been widely reported in bacterial and eukaryotic cells and can be the consequence of technical limits to the detection of low abundance mRNAs or proteins as well as the involvement of additional layers of post-transcriptional regulation (59-62). In the case of CdrS, its influence on proteasome subunit transcription could have knock-on influences in the proteome beyond transcriptional control. Our results are consistent with a hypothesis that there is substantial post-transcriptional regulation in the regulatory systems involving CdrS, and this may extend to other regulatory pathways in *H. volcanii*.

In Haloarchaea, the function of CdrS appears to be conserved in relation to the regulation of cell division (56). Our combined results suggest that CdrS is part of a regulatory network and controls the cell division apparatus and other downstream gene products in response to several conditions or stresses and via other transcription regulators. CdrS-mediated regulation of division might thereby play an important role in maintaining archaeal cell size homeostasis in coordination with metabolism, or by triggering changes in cell size or morphology in response to conditions or stress.

## Materials and Methods

### Strains and growth conditions

Strains, plasmids and oligonucleotides used are listed in Supplementary Table 4. *E. coli* strains DH5α (Invitrogen, Thermo Fischer Scientific, Waltham, USA) and GM121 (49) were grown aerobically at 37 °C in 2YT medium (Miller, J. H.,1972).

*H. volcanii* strains HV30 and HV35 (45) with plasmids were grown aerobically at 45 °C and 200 rpm in Hv-min medium or Hv-Ca (63, 64), or in media modified to contain an expanded trace elements and vitamin solution, which we refer to as Hv-MinTE (this study) or Hv-Cab medium (15, 16). Where necessary media was supplemented with uracil (10 µg/ml or 50 µg/ml as indicated) (for Δ*pyrE2*), and leucine (50 µg/ml) (for Δ*leuB*), and L-tryptophan with the indicated concentration (for Δ*trpA*). Tryptophan was also added to media to control gene expression of the *cas* genes coding for proteins Cas5-8b via the tryptophan inducible promoter (p.*tnaA*). Cultures were generally maintained in steady logarithmic growth (OD_650_ < 0.8) for at least 2 days prior to sampling (OD_650_ =0.2-0.8) for analysis, unless otherwise indicated. Data displayed are representative of at least three biological replicate experiments.

### Strains and plasmid construction

#### Generation of strain HV35

To generate a strain with inducible *cas*-gene expression, the gene cluster *cas6b, cas8b, cas7* and *cas5* was cloned downstream of the tryptophan inducible promoter p.*tnaA* and upstream of the terminator t.syn into the pTA131 plasmid that contained the up- and downstream regions of CRISPR locus C yielding pTA131-Cup-p.tnaA-cas6b8b75-t.syn-Cdo. Strain HV32 (44) was transformed with the plasmid pTA131-Cup-p.tnaA-cas6b8b75-t.syn-Cdo to mediate integration of the plasmid into the genome replacing CRISPR locus C. Pop-in candidates were plated on medium containing 5-FOA (5-Fluoroorotic acid) to select for pop-out clones. *cas*-gene insertion candidates were verified by Southern blot analysis (Supplementary Figure 9). Ten micrograms of genomic DNA were digested with *Sac*II and fragments were separated on a 0.8 % agarose gel. DNA fragments were transferred to a nylon membrane (Hybond™-N+) (GE Healthcare, Dornstadt, Germany) by capillary blotting. Two PCR products (termed Cup and cas8) with sizes of 305 bp (Cup) and 399 bp (cas8) were used as hybridisation probes. Fragment Cup was amplified using primers CdelupKpnI and CdelupiEcoRV and fragment cas8 was amplified using primers 5-HindIII-Cas8 and 8R126A#2. Probes were labeled using [α-^32^P]-dCTP and the DECAprime II DNA labeling kit (Thermo Fisher Scientific). Both hybridization and detection of the membranes were carried out as described in the manufacturer’s protocol. The strain resulting from this *cas*-gene integration with tryptophan inducible promoter p.*tnaA* was termed HV35.

#### Attempt to generate a *cdrS* deletion strain

To delete the *cdrS* gene the gene was amplified with 500 bp flanking regions by using primers HVO_0582-UP and HVO_0582_DO. The resulting PCR product was ligated into pTA131 (digested with *EcoR*V) resulting in plasmid pTA131-up-HVO_0582-do. Inverse PCR on pTA131-up-HVO_0582-do with primers iPCR_HVO 0582_UP and iPCR_HVO_0582_DO_NEU deleted the gene HVO_0582. After ligation of the resulting PCR product the plasmid pTA131-up-ΔHVO_0582-do was obtained. The wild-type strain H119 was transformed with pTA131-up-ΔHVO_0582-do to generate a knock-out strain with the pop-in/pop-out method (Bitan-Bitain). Pop-in clones were obtained and confirmed via colony-PCR with primers HVO_0582_UP and HVO_0582_DO. To obtain a pop-out strain 409 pop-in clones were screened with PCR with primers HVO_0582_UP and HVO_0582_DO. All clones still contained the HVO_0582 gene, suggesting that HVO_0582 is essential.

#### Plasmid pTA231-pfdx-HVO_0582NFlag used for ChIP-Sequencing

Primers 5’-HVO_0582-*Hind*III and 3’-HVO_0582-*Xba*I were used for amplification of the gene *cdrS* (HVO_0582) using genomic DNA from *H. volcanii* strain H119. The obtained DNA fragment was digested with *Hind*III and *Xba*I and ligated into pTA927 (digested with *Hind*III and *Xba*I) to yield pTA927-ptna-HVO_0582NFlag. This plasmid was digested with *Nde*I and *Xba*I and the resulting fragment was ligated into pTA231-pfdx (digested with *Nde*I and *Xba*I) resulting in pTA231-pfdx-HVO_0582NFlag.

#### Plasmids pTA232-tele-anti#1-3

Plasmids expressing the crRNAs from a t-element containing precursor were generated by inverse PCR with pMA-telecrRNA (45, 65) as template. The used primers (anti#1 fw, anti#1 rev/ anti#2 fw, anti#2 rev/ anti#3 fw, anti#3 rev) replace the spacer 1 of locus C with spacers anti#1, anti#2 or anti#3 against HVO_0582. Plasmids pMA-tele-anti#1/2/3 comprise the crRNA gene (8 nucleotide 5’
s handle/spacer/ 22 nucleotide long 3’ handle) flanked by t-elements. Plasmids were digested with *Kpn*I and *BamH*I and the resulting fragment was cloned into pTA232 (63) digested with the same enzymes, resulting in plasmids pTA232-tele-anti#1/2/3.

#### Plasmids pTA232-0582anti#1-3

Plasmids expressing the crRNA with flanking repeats were ordered (GeneArt, Thermo Fisher Scientific) as pMK-RQ-0582anti#1-anti#3. They contain a synthetic CRISPR locus, which comprises the leader of locus C, one spacer flanked by repeats and a synthetic terminator. After digesting the plasmids with *BamH*I and *Kpn*I to excise the entire loci, purified inserts were ligated into pTA232 digested with the same enzymes, yielding plasmids pTA232-0582anti#1-anti#3.

#### Construction of the complementation plasmids

Plasmids for complementation were generated via amplification of the genes HVO_0581, HVO_0582 or of both HVO_0582-HVO_0581 including the natural terminator with the primers (5’-HVO_0582-*Nde*I, 3’-HVO_0582-*Hind*III, 5’-HVO_0581-*Nde*I, 3’-HVO_0581-nat.t.-*Apa*I, 5’-*Hind*III-nat.t.). The PCR fragments were ligated into pBlue (that was digested with *EcoRV*) and resulting plasmids were digested with *Nde*I and *Apa*I to isolate the inserts that were ligated into plasmid pTA409-pfdx (digested with the same enzymes) to obtain the complementation plasmids pTA409-fdx-HVO_0581-nat.t, pTA409-fdx-HVO_0582-nat.t and pTA409-fdx-HVO_0582-HVO_0581-nat.t.

### Gene repression with CRISPRi

In *Haloferax*, two CRISPRi approaches that both work effectively can be employed (45). In one approach the crRNA gene is expressed between two t-elements that are processed by the cellular proteins RNase P and tRNase Z to release the mature crRNA. For this approach *Haloferax* strain HV30 is used. In a second approach the crRNA is expressed as part of a short synthetic CRISPR locus, that is processed by Cas6b to generate the mature crRNA; here *Haloferax* strain HV35 is used. For microscopic analyses of cell morphology and growth experiments, the crRNAs were encoded in a synthetic CRISPR locus and expressed in HV35. For all other experiments, the crRNAs were produced via the Cas6b-independent mechanism (65) in HV30. This strain is deleted for the *cas3* and *cas6b* genes, ensuring that DNA bound by Cascade is not degraded by Cas3 and endogenous crRNAs are not produced by Cas6b cleavage (Stachler and Marchfelder, 2016).

For repression of *cdrS* (HVO_0582), strain HV35 was transformed with the unmethylated (obtained via passage through *E. coli* GM121) CRISPR knock-down plasmid pTA232-0582-anti#1, pTA232-0582-anti#2, and pTA232-0582-anti#3 respectively, selecting the transformants on Hv-min agar medium with uracil (10 µg/ml or 50 µg/ml) and tryptophan (0.04 mM or 0.2 mM Trp) as indicated. Single colonies were streaked on the same medium, and colonies were screened by PCR to identify the presence of crRNA spacer (Supplementary Figure 10.) and named *cdrS* CRISPRi#1, *cdrS* CRISPRi#2, and *cdrS* CRISPRi#3. Since homologous recombination can happen between the repeats and thereby delete the crRNA spacer, it was important to test the strains with PCR to confirm the presence of the complete crRNA gene in the plasmids. For the complementation test, the strain HV35 was co-transformed with the unmethylated CRISPR knock-down plasmid (pTA232-0582-anti#1, pTA232-0582-anti#2, or pTA232-0582-anti#3) and the unmethylated expression plasmid (pTA409 as control, pTA409-fdx-HVO_0581-nat.t, pTA409-fdx-HVO_0582-nat.t and pTA409-fdx-HVO_0582-HVO_0581-nat.t). Selection for transformants containing two plasmids was achieved by growth on Hv-MinTE agar medium with L-tryptophan (0.04 mM Trp). Single colonies were streaked on the same medium, and colonies were screened by PCR to identify the presence of the gene for the crRNA spacer (Supplementary Figure 10).

For the overexpression test, the strain HV35 was transformed with the unmethylated expression plasmid pTA409 (as a control), pTA409-fdx-HVO_0581-nat.t, pTA409-fdx-HVO_0582-nat.t and pTA409-fdx-HVO_0582-HVO_0581-nat.t, followed by selecting the transformants on Hv-Cab agar medium with L-tryptophan (0.04 mM or 0.2 mM). The overexpression effect was also tested in *H. volcanii* strain H26 (Δ*pyrE2*) by selecting the transformants on Hv-Cab agar medium.

### Growth experiments

Cells were grown in Hv-MinTE medium with addition of tryptophan (0.08 mM) and uracil (50 µg/ml) aerobically with shaking (200 rpm) at 45°C. Cell growth was monitored by measurement of the optical density at 650 nm wavelength (OD_650_). For each strain, three biological replicates were prepared. Doubling times d for strains were calculated as follows: the natural logarithm of 2 divided by the growth rate µ (d = ln 2/µ). The growth rate µ itself is calculated through the natural logarithm of the values of the time range divided by the time range (µ = (ln x(t)-ln x(t_0_)) / (t-t_0_)).

### Light Microscopy

For most phase-contrast microscopy, a two microliter sample of culture was placed on an agarose pad prepared by dropping ∼ 50 µl of 1% agarose containing 18% BSW (includes calcium) (64) onto a glass slide at room temperature, and a clean 22 × 50 mm number 1.5 glass coverslip placed on top. Images were acquired using a 100 × / 1.4 NA oil immersion objective and phase contrast optics using a Zeiss Axioplan microscope (Carl Zeiss, Oberkochen, Germany).

### Coulter cell volume analysis

Culture samples were diluted (1:100 or 1:1000) with 0.2-micron filtered 18% BSW, and were analysed with a Multisizer 4 Coulter Counter (Beckman Coulter, Brea, USA) equipped with a 30 nm aperture tube, calibrating with 2-micron latex bead standard (Beckman Coulter) diluted in 18% BSW as the electrolyte. Runs were completed in volumetric mode (100 μl) and current set to 600 μA and gain set to 4.

### Cell shape quantification

Microscope images were analysed using ImageJ 1.53c. Phase-contrast images were first smoothed using a Gaussian filter followed by a Rolling-ball background subtraction. Individual objects (cells) were then identified by optimized thresholding. Parameters for detection were adjusted and touched cells were manually split. The holes in the binary objects were filled by using a hole filling operation. Cell shape parameters were determined by using ‘Analyze particles’ function. The minimum cell size was 0.2 μm^2^ and the edged objects were excluded. The circularity of each cell was calculated as the percentage of cell area to the minimal circle area that completely contains the cell outline (15), by using a custom macro implemented in ImageJ (16). The cell division mutants were quantified as four distinct cell shapes: filaments (cell area >= 6.5 μm^2^, circularity <=0.4), giant plate cells (cell area >= 6.5 μm^2^, circularity > 0.4); cellular debris (cell area <= 2 μm^2^), wild-type-like cells (cell area between 2 μm^2^ and 6.5 μm^2^).

### Northern Blots analysis

#### For CRISPRi strains

RNA was isolated from strains HV30 x pTA232-tele-0582anti#1-3 and wild-type strain HV30 x pTA232 as described before (66). Ten micrograms of RNA were separated on a denaturing 8% polyacrylamide gel or a denaturing 1% agarose gel and transferred to a nylon membrane (Hybond N^+^ membrane or Pall membrane). For hybridisation experiments radioactively labeled PCR probes against the desired targets were generated using DECAprime II DNA labeling kit and [α-^32^P]dCTP (Life Technologies, Thermo Fisher Scientific). Membranes were subsequently incubated with the labeled PCR fragments.

For the quantification of the repression efficiency northern membranes were exposed to imaging plates and analyzed with the Amersham Typhoon Biomolecular Imager and the ImageQuantTL™ software. Signals were compared with the signals for the 16S rRNA used as a loading control. The amount of the detected RNA signals for the wild-type controls was set to 100%. Northern blot analyses were done in triplicates.

#### For complemented strains

The *cdrS* CRISPRi strain (expressing crRNA#2) was complemented with either the single gene HVO_0581 (*ftsZ2*) or with the complete operon HVO_0582-HVO_0581. Transformed strains were grown in minimal medium supplemented with tryptophan (0.25 mM) and harvested in exponential phase (OD_650nm_ ≈ 0.4-0.52). RNA was isolated as described before (66), separated on a 1% agarose gel and transferred onto a nylon membrane (Pall membrane). For hybridisation experiments radioactively labeled PCR probes against the selected targets were generated using DECAprime II DNA labeling kit and [α-^32^P]dCTP (Life Technologies, Thermo Fisher Scientific). Membranes were subsequently incubated with the labeled PCR fragments against the mRNA of the cluster HVO_B0192-HVO_B0193 and against the mRNA of HVO_0739.

### ChIP-Sequencing

#### Preparation of the samples

The ChIP-Seq analysis was performed with the FLAG-tagged CdrS and a control with three replicates each. The culture HV35 x pTA231-pfdx-HVO_0582NFlag was grown in Hv-Ca medium supplemented with uracil (50 µg/ml) to an OD_650nm_ of 0.7-0.85 (replicate1: 0.74, replicate2: 0.72, replicate3: 0.85). Crosslinking was done with a final concentration of 0.5% (v/v) formaldehyde for 20 min at room temperature under constant shaking. After 20 minutes the reaction was stopped by addition of glycine to a concentration of 0.25 M. Cells were harvested at 4°C and 9,800 x g for 20 min and washed twice with enriched phosphate buffer (PBS). Cells were resuspended in the appropriate volume of lysis puffer (V_culture_ x OD_650nm_ / 45 = ml lysis puffer). After ultrasonification using a Branson Sonifier, the solution was centrifuged with 15,500 x g at 4 °C for 1 hour yielding the S15 protein fraction. The DNA was fragmented to an average length of 200-500 bp with ultrasonification for 60 min with a duty cycle of 50%, followed by a RNA digestion with RNase A for 30 min at 37 °C. For the control, 10% of the S15 extract was removed and later processed in the same way as the purified protein-DNA complex was. The FLAG-tagged protein-DNA complex was purified via affinity chromatography using anti-FLAG M2 affinity gel (Sigma-Aldrich, Taufkirchen, Germany). The purified protein-DNA complexes and the control were incubated at 95 °C for 40 min for reversal of the crosslink. DNA samples were subsequently incubated with RNase A and proteinase K for 20 min at 37 °C. After a phenol-chloroform extraction DNA was precipitated and resolved in 10 μl DNase-free water, concentration was measured with a NanoDrop photometer.

#### Library preparation and sequencing

DNA library preparation was carried out according to the manufacturer’
ss protocol using the NEXTflex™ ChIP-Seq Kit (NOVA-5143-01) (PerkinElmer, Hamburg, Germany) and the NEXTflex™ ChIP-Seq Barcodes (NOVA-514122) (PerkinElmer). DNA libraries were pooled to 4 nM and sequenced using an Illumina MiSeq.

#### Bioinformatics analyses

In order to remove sequencing errors, a quality control was applied to the sequenced reads in FASTQ files using FastQC tool (21), then we used the TrimGalore tool (https://www.bioinformatics.babraham.ac.uk/projects/trim_galore/) to remove adapters and perform read quality trimming. The reads were then aligned to the main chromosome and the three plasmids pHV1,3 and 4 via Bowtie2 (67). After that, we computed the correlation between read counts in different regions for all samples using deeptool (68). Next, we identified enriched binding sites (peaks) using MACS2 callpeak (69). Finally, MEME-ChIP (48) tool was used for discovering motifs in the peaks regions.

### Transcriptome analyses

Strain HV30 x pTA232-tele-anti#2 and HV30 x pTA232 were grown in Hv-MinTE medium supplemented with tryptophan (0.25 mM) and uracil (50 µg/ml) to exponential phase (OD_650nm_=0.42-0.47), experiments were done in triplicates. RNA was isolated as described for northern analyses and sent to Vertis Biotechnologie AG (Munich, Germany) for cDNA synthesis and sequencing. From the total RNA samples, ribosomal RNA molecules were depleted using the Ribo-Zero rRNA Removal Kit for bacteria (Illumina, San Diego, USA). The ribodepleted RNA samples were first fragmented using ultrasound (4 pulses of 30 s each at 4°C). Then, an oligonucleotide adapter was ligated to the 3’ end of the RNA molecules. First-strand cDNA synthesis was performed using M-MLV reverse transcriptase and the 3’ adapter as primer. The first-strand cDNA was purified and the 5’ Illumina TruSeq sequencing adapter was ligated to the 3’ end of the antisense cDNA. The resulting cDNA was PCR-amplified to about 10-20 ng/μl using a high-fidelity DNA polymerase. The cDNA was purified using the Agencourt AMPure XP kit (Beckman Coulter). For Illumina sequencing, the cDNAs were pooled in approximately equimolar amounts. The library pool was fractionated in the size range of 200 - 550 bp using a differential clean-up with the Agencourt AMPure kit. The cDNA pool was sequenced on an Illumina NextSeq 500 system using 75 bp read length.

#### Bioinformatics analyses

A quality control was applied to the sequenced reads in FASTQ files using FastQC tool (21), then TrimGalore tool (https://www.bioinformatics.babraham.ac.uk/projects/trim_galore/) was used to remove adapters and perform read quality trimming. For analysis of transcriptome samples, reads were aligned to the main chromosome and mini-chromosomes (pHV1, pHV3, and pHV4) using Bowtie2 (67). Differential gene expression analysis was performed using the DESeq2 (70) with default settings. We considered the genes with an adjusted p-value less than or equal to 0.005 as significantly differentially expressed. With this cut-off, we allowed less than 1% of the false positives from the significantly differentially expressed genes.

### Proteome analyses

Strains were grown as described for transcriptome analyses. After centrifugation cells were resuspended in 18% salt water supplemented with 0.1 mM PMSF, 1 mM benzamidine, 1 µg/ml pepstatin A, 1µg/ml leupeptin and 10 mM DTT. After ultrasonification with a 50% duty cycle for 3 min, taurodeoxycholate was added to a final concentration of 0.006%. The suspension was subsequently incubated for 30 min at 4 °C, followed by ultracentrifugation at 30.000 x g for 45 min at 4 °C. The supernatant and the pellet (which was resuspended with 50 mM Tris-HCl puffer (pH7)), were incubated with 20 µl DNaseI (RQ1), 0.2 µl Exonuclease III and 0.5 µl RNaseA for 1 h at 4 °C with gentle shaking. Soluble proteins from the supernatant fraction were precipitated with acetone. Insoluble proteins were purified with StrataClean beads (Agilent, Santa Clara, USA).

For analysis of the soluble protein fraction 50 µg protein were reduced with 2.5 mM TCEP (tris-(2-carboxyethyl) phosphine hydrochloride) (Invitrogen, Thermo Fisher Scientific) at 65 °C for 45 min before thiols were alkylated in 5 mM iodoacetamide (Sigma-Aldrich) for 15 min at 25 °C in the dark. For protein digestion trypsin (Promega, Walldorf, Germany) was added in an enzyme to substrate ratio of 1:100 before incubation at 37 °C for 14 h. For digestion of proteins in the insoluble fraction 20 µl StrataClean beads (Agilent), which bound 20 µg protein, were incubated with 2.5 mM TCEP at 65 °C for 45 min before 5 mM iodoacetamide was added. Samples were incubated for 15 min at 25 °C in the dark and subsequently digested with trypsin (enzyme to substrate ratio 1:100) for 14 h at 37 °C. To ensure the complete generation and removal of peptides from the beads, further trypsin (200 ng) was added and incubation at 37 °C was prolonged for additional 3 h. Peptides were eluted from the beads by incubation in a sonication bath for 2 min and transfer of the supernatant to a new vial. The beads were than washed sequentially with 100 µl solvent A (0.1% acetic acid in water) and 100 µl 60% solvent B (0.1% acetic acid in acetonitrile diluted in solvent A). After each washing step peptides were again eluted by sonication and supernatants were collected in the same vial as the initial elution. Subsequently peptides were dried by evaporation of the solvents and suspended in 10 µl solvent A before MS analysis.

For MS analysis peptides were loaded on an EASY-nLC 1000 system (Thermo Fisher Scientific) equipped with an in-house built 20 cm column (inner diameter 100 µm, outer diameter 360 µm) filled with ReproSil-Pur 120 C18-AQ reversed-phase material (3 µm particles, Dr. Maisch GmbH, Ammerbuch-Entringen, Germany). Elution of peptides was executed with a nonlinear 180 min gradient from 1 to 99% solvent B (0.1% (v/v) acetic acid in acetonitrile) with a flow rate of 300 nl/min and injected online into an Orbitrap Q Exactive (Thermo Fisher Scientific) in DIA mode. In each DIA cycle one survey scan at a resolution of R = 70,000 at m/z 200, 120 ms maximal IT, 3×10^6^ automatic gain control target and mass range 300-1250 followed by 22 variable width DIA scans at a resolution of R = 35,000 at m/z 200, automatically set maximal IT, 2×10^5^ automatic gain control target and a NCE of 27.5 were obtained. All scans were acquired in the Orbitrap with activated lock mass correction. The cycle time was 3.6 s ensuring at least 8 scans over the width of a typical LC peak (30 seconds).

Data analysis was performed using Skyline (version 20.1.0.31)(71) applying an in-house built spectral library of peptides identified from previous in-house analysis of *H. volcanii*. Exported transition peak areas were converted to protein quantities using the MaxLFQ algorithm implemented in the iq R package (72).

Proteins that were quantified in all three biological replicates of either the CRISPRi strain or the wild-type strain but in none replicate of the other strain were considered as on (only in CRISPRi strain) or off (only in wild-type strain). Proteins which were detected in at least two biological replicates of each strain were included for relative quantification of protein abundance. Data were considered statistically significant when -log_10_(p-value)>2. Biological significance was assumed for quantified proteins exhibiting a log_2_ fold change in abundance>|2| when comparing CRISPRi and wild-type strain.

All MS data (raw data, data analysis results, spectral library) have been deposited to the ProteomeXchange Consortium via the PRIDE partner repository (73) with the dataset identifier PXD017903.

### Western blotting

The protein expression levels of FtsZ1, and FtsZ2 were assessed by western blot using rabbit antisera for FtsZ1 and FtsZ2 (15). FtsZ1 and FtsZ2 rabbit antisera were generated with a synthetic peptide antigen derived from the sequence of the C-terminal region of FtsZ1: [C]-Ahx-QAHAEERLEDIDYVE-acid (Cambridge Research Biochemicals, Billingham, UK) and C-terminal region of FtsZ2: NH2-[C]-SDGGRDEVEKNNGLDVIR-COOH (Thermo Fisher Scientific). *H. volcanii* cell pellets were resuspended in SDS–PAGE sample buffer and then heated (95 °C, 5 min) and vortexed. Same amount of protein was separated by SDS-PAGE, electroblotted (Bio-Rad, Feldkirchen, Germany) onto a nitrocellulose membrane (Protran) (Sigma-Aldrich), probed with rabbit polyclonal primary antiserum (1:1000 dilution for FtsZ1, or 1:2000 dilution for FtsZ2) followed by a secondary antibody of donkey anti-rabbit-IgG (AbCam16284, 1:5000 dilution) conjugated to horseradish peroxidase. Protein bands were detected using an enhanced chemiluminescence detection kit (Thermo Fisher Scientific) and visualized and quantified by Amersham imager 600 instrument (GE Healthcare). The western blot signals were quantified using Image J 1.53c software. The relative grey intensities for FtsZ1 and FtsZ2 protein bands were normalized to the lane’s loading control (with Ponceau S staining). All western blots were performed at least twice with independent biological samples and showed similar results. The western blot results are shown for one experiment, but quantification was from at least two independent experiments.

## Acknowledgments

We thank Susanne Schmidt for expert technical assistance. Funding was provided as follows: by the Australian Research Council (FT160100010) to Iain G. Duggin, by DFG priority programs SPP2002 (BA 2168/21-1) and SPP 2141 (BA 2168/23-1) to Rolf Backofen, by DFG priority program SPP2002 (BE 3869/5-1) to Dörte Becher and by DFG priority program SPP2002 (DFG Ma1538/24-1) to Anita Marchfelder.

## Competing interests

The authors do not have any competing interests.

## Footnotes

1 We define small proteins here as proteins of 70 or less aa.

2 We term the HV30 and HV35 strains here as wild-type strains and the HV30 and HV35 strains that express the crRNAs as CRISPRi strains.

## Notes

### Competing Interest Statement

The authors have declared no competing interest.

