## Supplementary Data for "CdrS is a global transcriptional regulator influencing cell division in *Haloferax volcanii*"

### Contents:

Supplementary Figures 1-10

page:

Supp. Figure 1. CRISPRi system for repression of *cdrS* 2
Supp. Figure 2. Quantification of *cdrS* and *cdrS* and *ftsZ2* repression 3
Supp. Figure 3. ChIP-Seq between the genes *hisB* (HVO_2986) and HVO_2987 4
Supp. Figure 4. Cell shape quantification analysis of *cdrS* repression 5
Supp. Figure 5. Cell division defects at different conditions 6
Supp. Figure 6. Complementation expression CRISPRi cells. 7
Supp. Figure 7. Overproduction of FtsZ2 and/or CdrS in HV35 background 8
Supp. Figure 8. Cell shape quantification analysis of *cdrS* overexpression 9
Supp. Figure 9. Southern blot analysis of strain HV35 10
Supp. Figure 10. PCR analysis to confirm the presence of *cdrS* targeting spacer 11

Supplementary Tables 1-6

page:

Supp. Table 1. Genes up- and downregulated in *cdrS* CRISPRi cells. 12
Supp. Table 2. Differentially abundant proteins in wild-type vs CRISPRi cells. 15
Supp. Table 3. Quantitative analysis of cell shape. 17
Supp. Table 4. Strains, plasmids and oligonucleotides used 18
Supp. Table 5. Complete list of up- and down regulated genes, separate Excel file.
Supp. Table 6. Complete list of protein abundances, separate Excel file.

References

page: 22

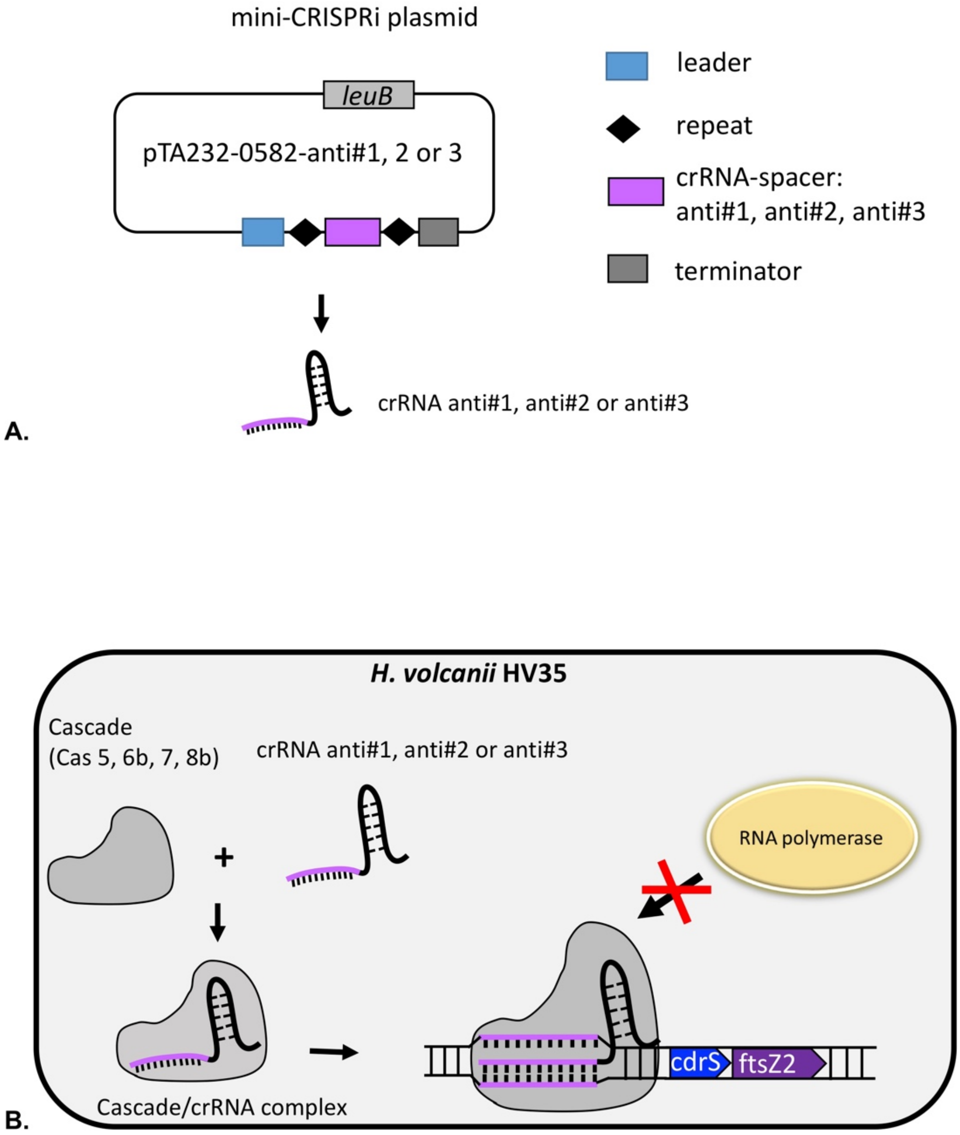


**Supplementary Figure 1. CRISPRi inhibits transcription initiation.**
**A.** The mini-CRISPRi plasmid used for expression of crRNAs. The plasmid contains the CRISPR leader sequence which contains the promoter, one crRNA spacer flanked by two repeats, and a synthetic terminator. The crRNA is expressed from the leader promoter and terminated by a synthetic terminator. The *leuB* gene is used as marker gene for selection in *H. volcanii.* **B.** Schematic illustration of CRISPRi targeting of the *cdrS* promoter region in *H. volcanii*. Cascade (complex of Cas proteins Cas 5, 6b, 7, and 8b) binds crRNA anti#1, anti#2, or anti#3 expressed from the plasmid (panel A) to form the Cascade/crRNA complex. The Cascade/crRNA complex is guided by the crRNA to bind to the target DNA sequence at the promoter and TSS region of *cdrS*, thereby preventing transcription initiation.

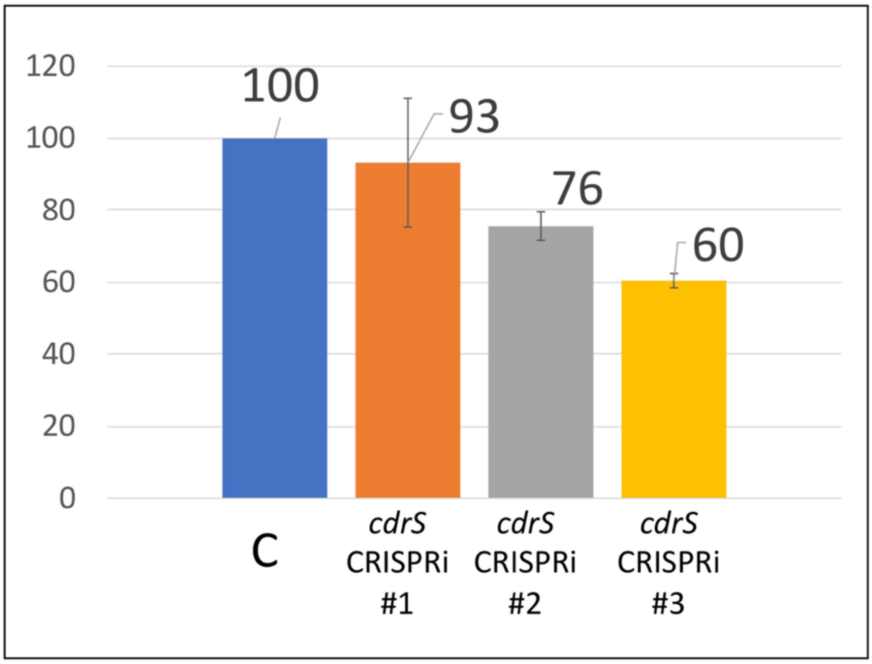


**A.**

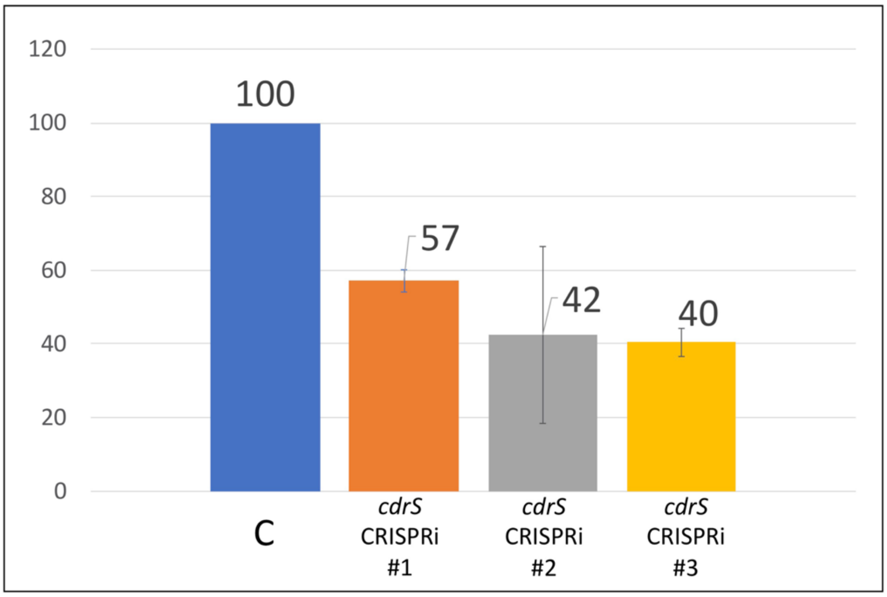


**B.**

**Supplementary Figure 2. Quantification of *cdrS* and *cdrS-ftsZ2* repression.** Signal intensity of the the bicistronic *cdrS-ftsZ2* mRNA (panel A) and the *cdrS* mRNA (panel B and was measured with ImageJ and set into relation to the 16S rRNA signal that was used as loading control. The proportion of the detected RNA is given in percentage compared to cells carrying a control plasmid. The amount of RNA in the control strains was set to 100% (column C) and of those expressing a targeting spacer (the three CRCISPRi strains expressing anti#1-#3) were set in relation. The strongest repression of the *cdrS* mRNA is obtained upon expression of crRNA anti#3.

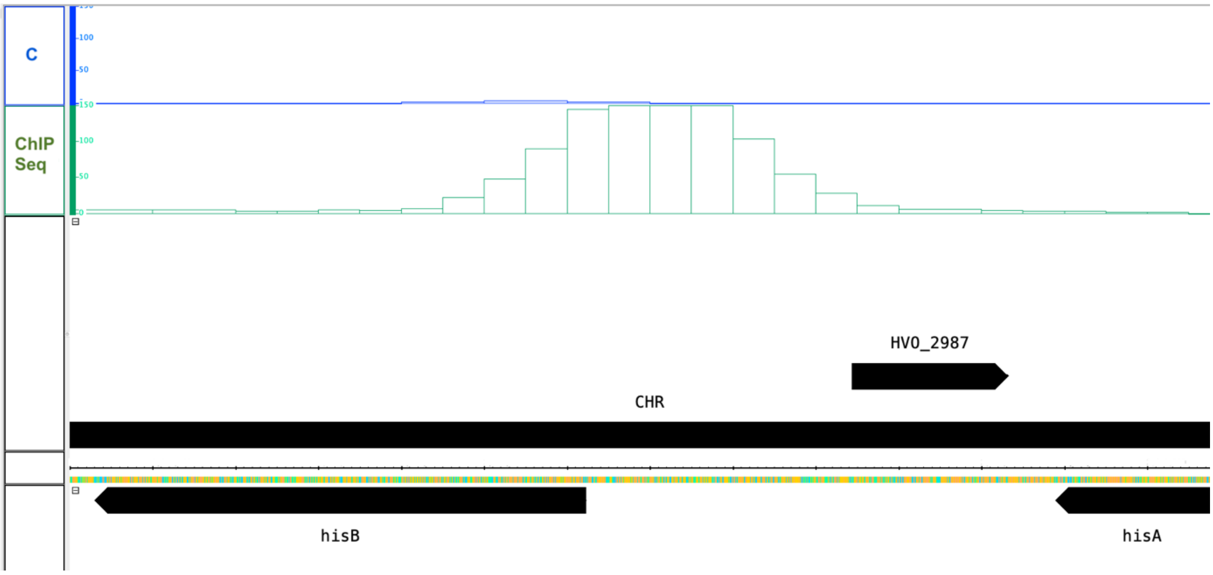


**Supplementary Figure 3. ChIP-seq data from the region between the genes *hisB* (HVO_2986) and HVO_2987.**

Here the CdrS binding motif is found on both strands, on the plus strand upstream of HVO_2987 and on the minus strand upstream of the *hisB* gene (HVO_2986). The control is shown in blue in the upper panel (panel C) and the ChIP-seq sample with CdrS is shown in green (panel ChIP-seq). Read numbers are indicated at the left.

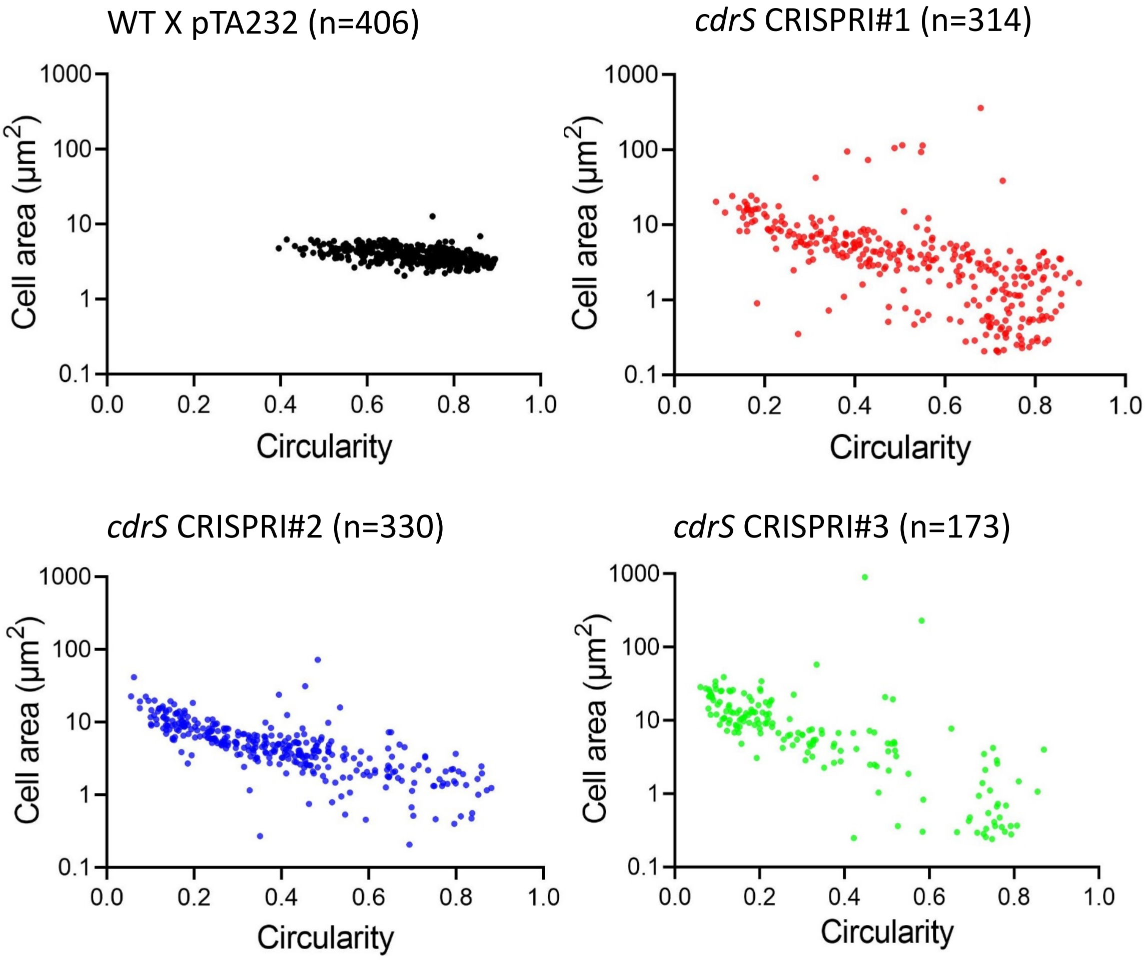


**Supplementary Figure 4. Cell shape quantification analysis of cdrS repression during mid-log growth in Hv-MinTE medium (+ 10 µg/ml uracil + 0.04 mM Trp).** Scatter plots of single cell values for cell area (µm$^2$) vs cell circularity for wild-type (HV35 × pTA232, n=406), *cdrS* CRISPRi#1 (HV35 × pTA232-0582anti#1, n=314), *cdrS* CRISPRi#2 (HV35 × pTA232-0582anti#2, n=330); and *cdrS* CRISPRi#3 (HV35 × pTA232-0582anti#3, n=173). The percentages of cells in the size and shape categories (see Methods) were: *cdrS* CRISPRi#1, filamentous 20%, giant-plate cell 5.7 %; *cdrS* CRISPRi#2, filamentous 30 %, giant-plate cell 3 %; *cdrS* CRISPRi#3, filamentous 52.6 %, giant-plate cell 4.6 %)

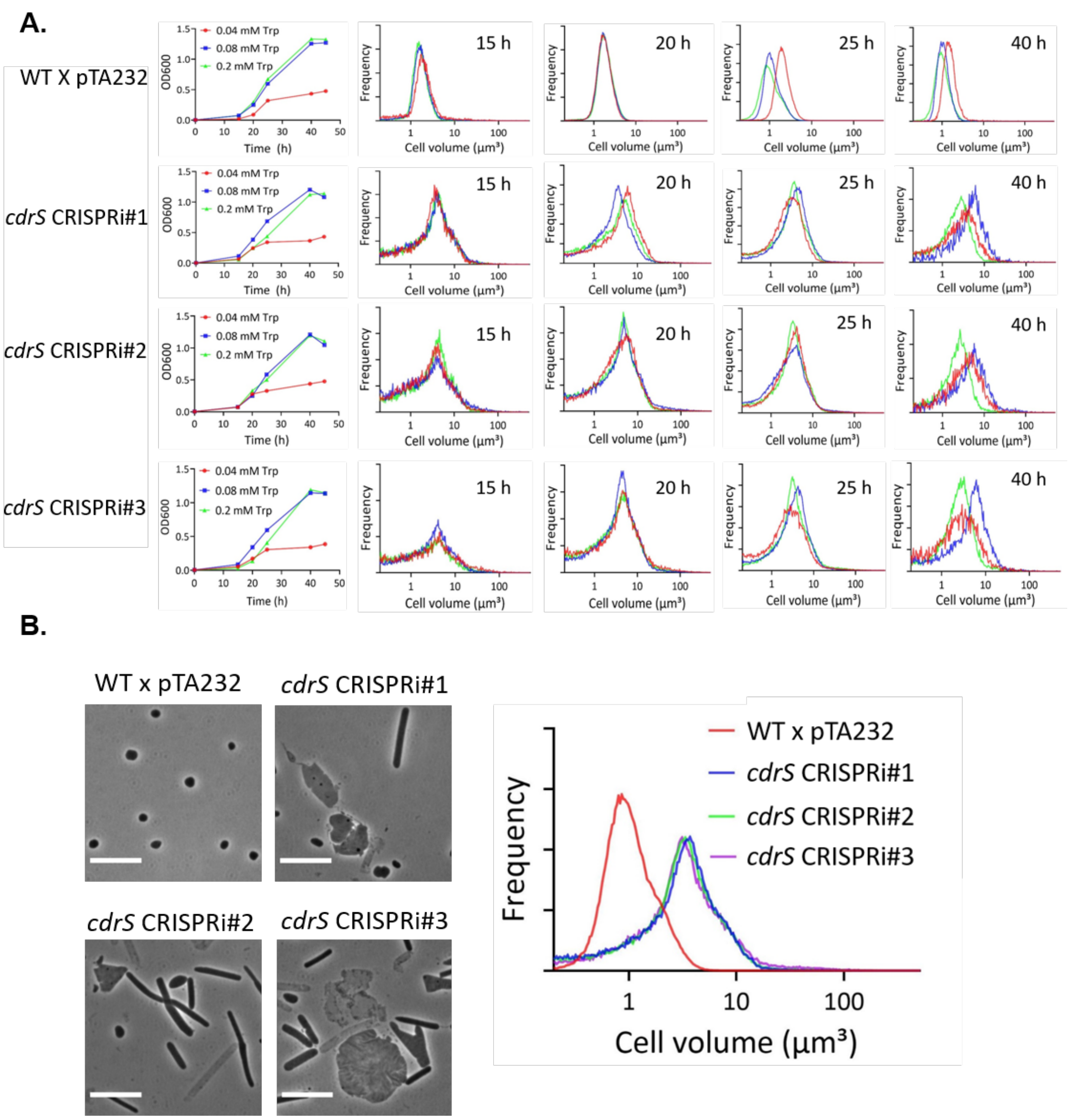


**Supplementary Figure 5. Different growth stages and tryptophan concentrations did not induce different effects of *cdrS* CRISPRi on cell division defects.**
**A.** Growth of four strains including wild-type (HV35 × pTA232), *cdrS* CRISPRi#1, *cdrS* CRISPR#2, and *cdrS* CRISPR#3 were performed in Hv-MinTE medium (+ 10 µg/ml uracil) under 0.04 mM, 0.08 mM, and 0.2 mM Trp (left vertical plane). Expression of the mRNAs for the Cascade complex proteins is regulated by the p.*tnaA* promoter, which is induced by addition of tryptophan, more tryptophan results in more Cascade complexes. The cell volume of four strains were analyzed under 0.04 mM Trp (red), 0.08 mM Trp (blue), and 0.2 mM Trp (green) at different growth time points 15 h, 20 h, 25 h, and 40 h. **B.** Phase-contrast images and Coulter cell volume analyses of wild-type, *cdrS* CRISPRi #1, *cdrS* CRISPRi#2, and *cdrS* CRISPRi#3 strains sampled at 25 h (log growth) under 0.2 mM Trp. All the three *cdrS* CRISPRi strains showed similar cell division defects compared to growth under 0.04 mM Trp (see Fig. 5). Scale bars, 10 µm. The cell volume of three CRISPRi strains showed similar and larger size distributions compared to the wild-type.

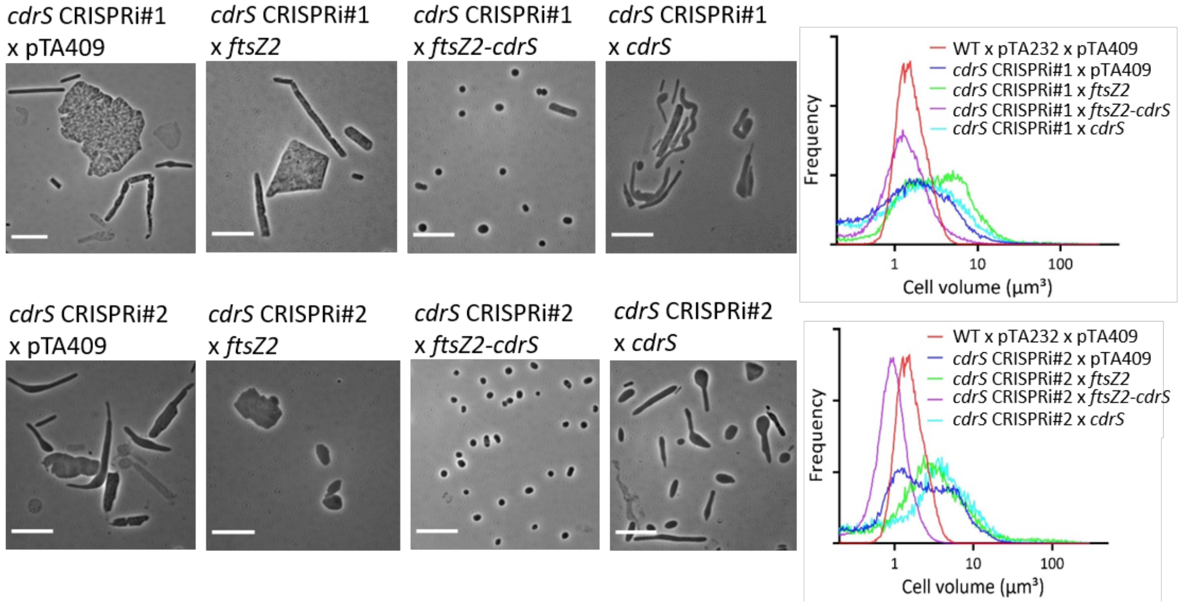


**Supplementary Figure 6. Characterization of cell division phenotype upon complementation expression of *ftsZ2*, *ftsZ2-cdrS*, and *cdrS* in *cdrS* CRISPRi#1 and *cdrS* CRISPRi#2 backgrounds.**

Both the phase-contrast images (right panel) and the corresponding Coulter cell volume analysis (left panel) of *cdrS* CRISPRi#1 complementation strains (upper) and *cdrS* CRISPRi#2 complementation strains (bottom) showed that the supplemental expression of both *ftsZ2* and *cdrS* restored cell division, whereas single complementation expression of *ftsZ2* or *cdrS* failed to rescue. All the strains were sampled during steady mid-log phase in Hv-MinTE medium supplemented with 0.04 mM Trp. Scale bars, 10 μm.

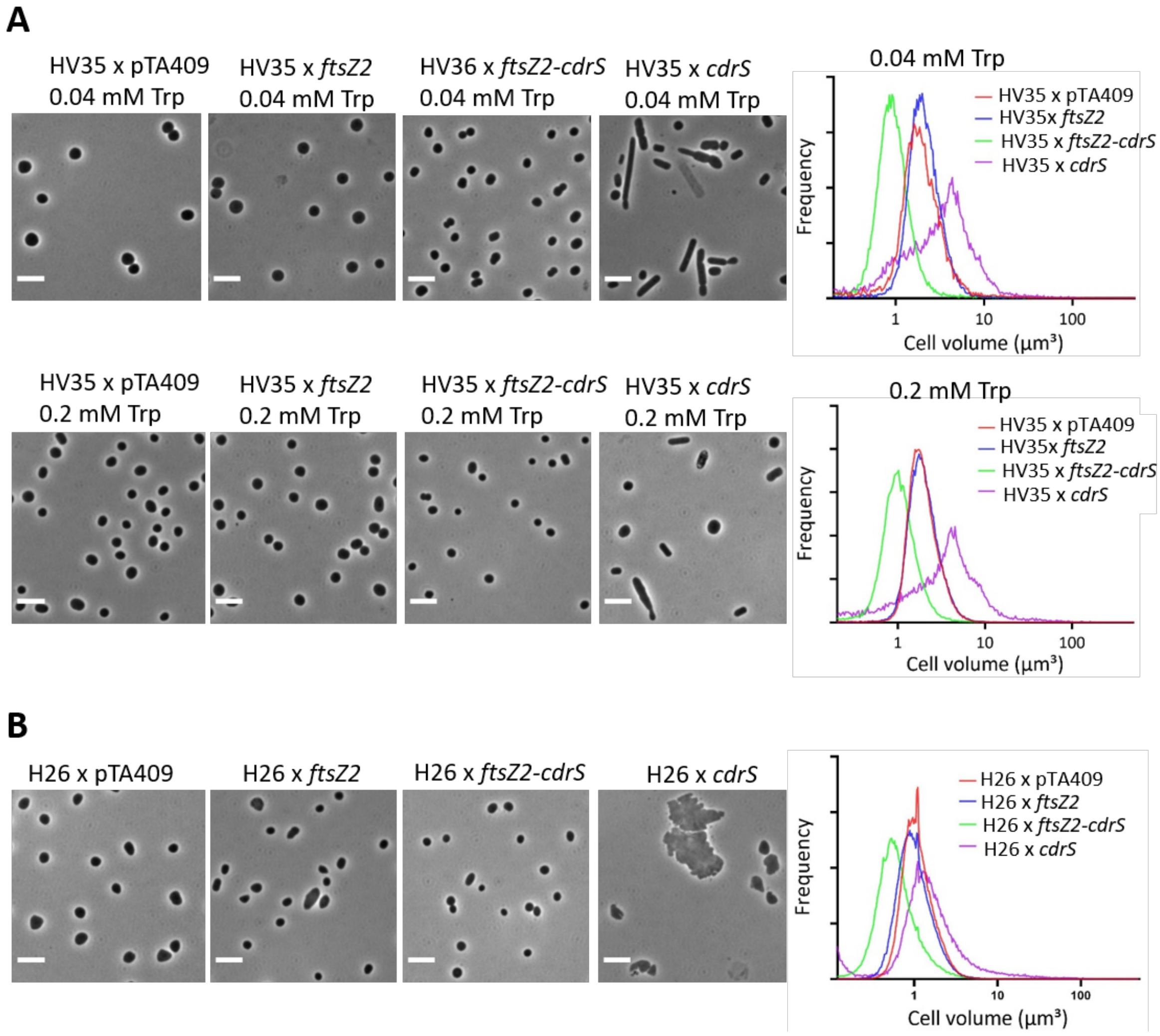


**Supplementary Figure 7. Characterization of cell division phenotype upon overproduction of FtsZ2 and/or CdrS in HV35 background with different concentration of Trp (A) and in H26 (Δ*pyrE2*) without Trp (B).**

**(A)** The phase-contrast images (right panel) and the corresponding coulter cell volume analysis (left panel) of wild-type control (HV35 x pTA409), *ftsZ2* overexpression (HV35 x *ftsZ2*), *ftsZ2-cdrS* overexpression (HV35 x *ftsZ2*), and *cdrS* overexpression (HV35 x *ftsZ2*) sampled in the Hv-MinTE medium (+ leucine 50 µg/ml) with 0.04 mM Trp (top panel) or 0.2 mM Trp (bottom panel). The data showed that double *ftsZ2-cdrS* overexpression resulted in wild-type morphological cells with significantly smaller cell size, while overexpression of single *cdrS* led to inefficient or mis-regulated division. Scale bars, 5 µm. **(B)** Trp-independent *ftsZ2* and/or *cdrS* overexpression effect on cell division in H26. Phase contrast micrographs (right) and corresponding coulter cell volume analysis of H26 control (H26 x pTA409), and *ftsZ2* overexpression, *ftsZ2-cdrS* overexpression, and *cdrS* overexpression in H26 background strains sampled during steady mid-log growth in Hv-Cab medium. The overexpression of *ftsZ2* and/or *cdrS* in H26 without Trp in the medium showed similar effect in HV35 background with different concentrations of Trp in the medium. Scale bars, 5 µm.

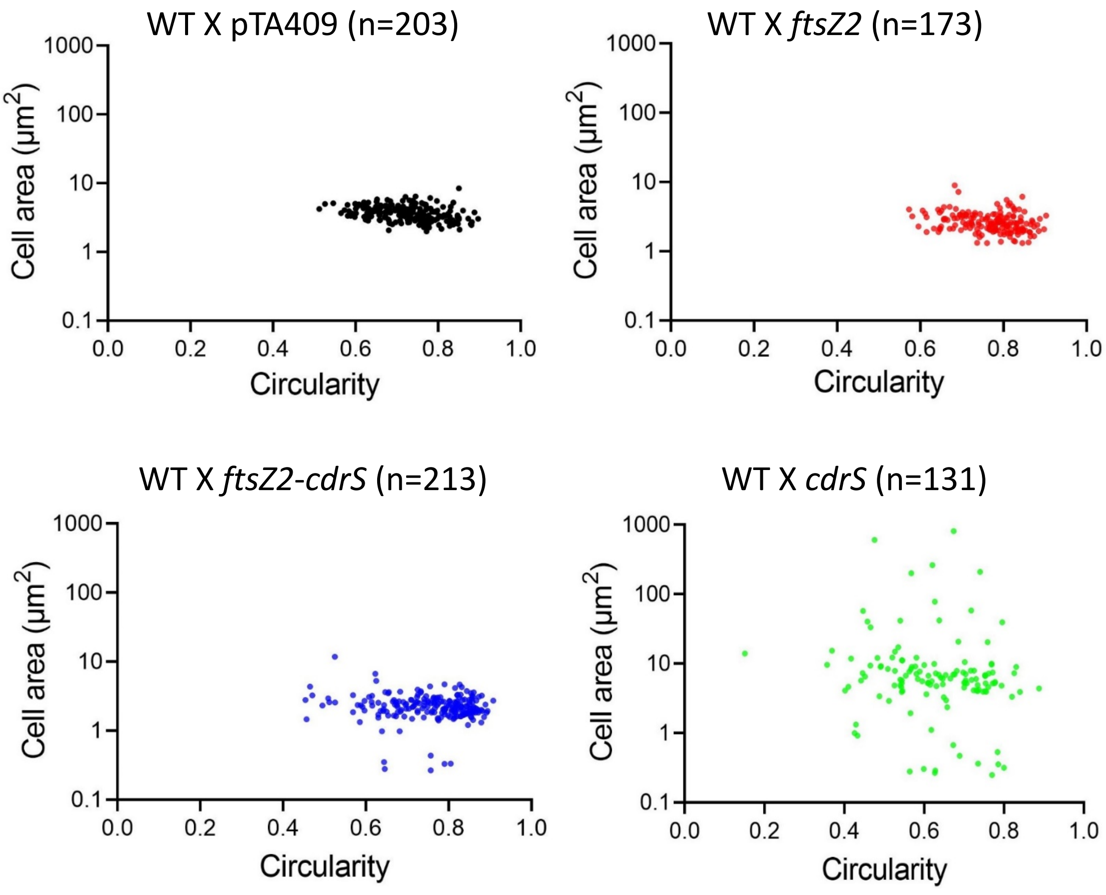


**Supplementary Figure 8. Cell shape quantification analysis upon overproduction of FtsZ2 and/or CdrS in wild-type background.**

Scatter plots of single cell values for cell area ($\mu m^2$) vs cell circularity for wild-type (HV35 × pTA409, n=203), FtsZ2 overexpression (HV35 × pTA409*ftsZ2*, n= 173), FtsZ2-CdrS double overexpression (HV35 × pTA409*ftsZ2-cdrS*, n=213), and CdrS overexpression (HV35 × pTA409*cdrS*, n=131). Samples were taken from mid-log growth in Hv-MinTE medium (+ leucine 50 µg/ml) with 0.08 mM Trp (Figure 7 sample).

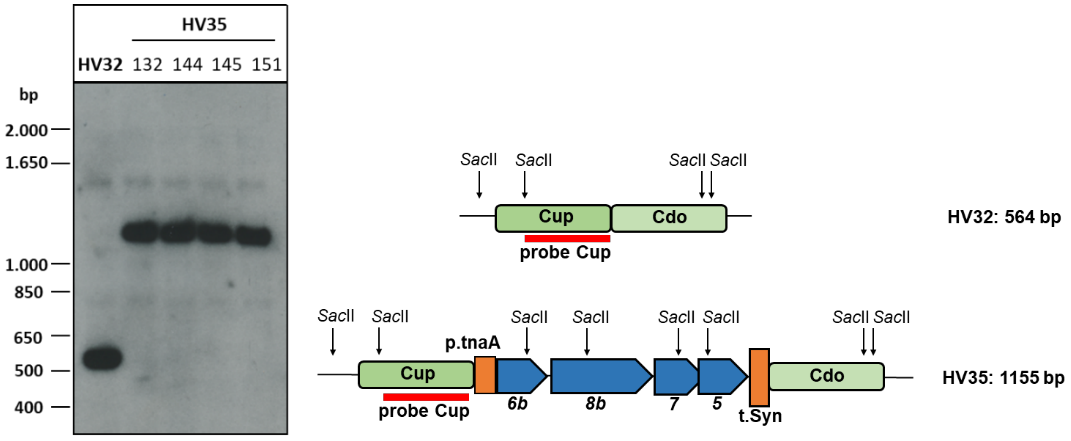


A.

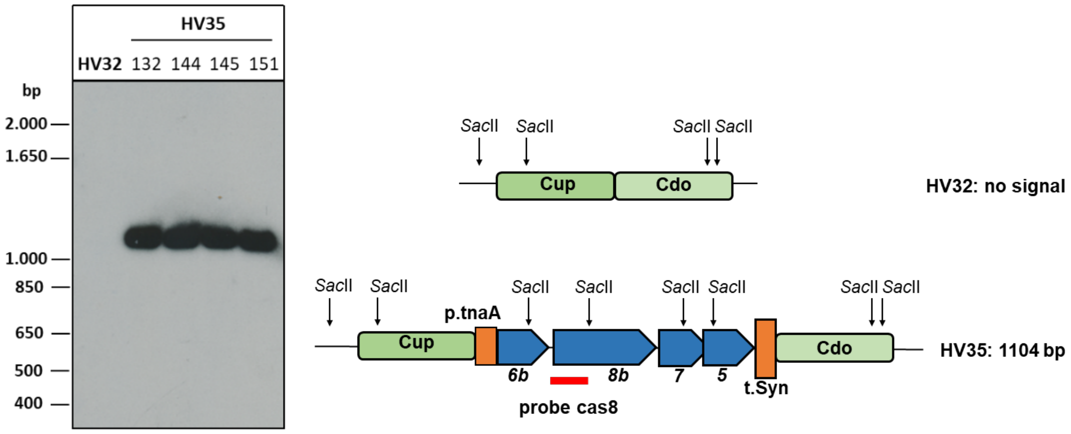


B.

**Supplementary Figure 9. Southern blot analysis of strain HV35.**
To verify the *cas*-gene insertion, genomic DNA from HV32 and four potential HV35 clones was isolated and 10 µg were digested by *Sac*II. After separation of DNA fragments on a 0.8% agarose gel, fragments were transferred by capillary blotting to a nylon membrane (Amersham Hybond™-N+, GE Healthcare) and fixed on the membrane by UV crosslinking. **A. Hybridisation with a probe against the upstream region.** Expected signals were 564 bp for HV32, in case of *cas*-gene integration in HV35, the expected size was 1,155 bp. **B. Hybridisation with a probe against *cas8b*.** Since the probe cas8 binds to the *cas8b* gene, no signal was expected for HV32, HV35 strains showed the expected signal of 1,104 bp. Genomic locations are shown schematically at the sides.

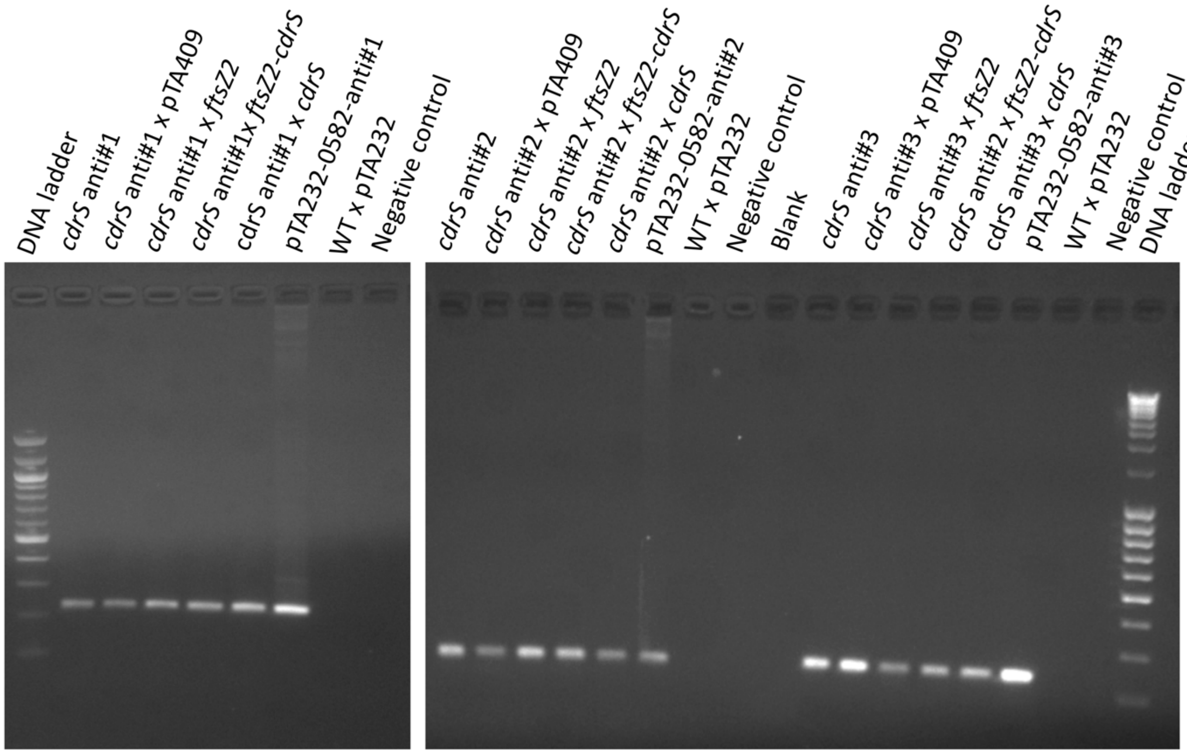


**Supplementary Figure 10. PCR analysis of all the *cdrS* CRISPRi and complementation strains to confirm the presence of *cdrS* targeting spacer.** The genomic DNA was extracted from the strains tested including wild-type vector only (HV35 x pTA232), all the three *cdrS* CRISPRi strains, and *ftsZ2*, *cdrS* and double *ftsZ2-cdrS* complementation strains in the corresponding *cdrS* CRISPRi strains, and used as the template for the PCR to verify the presence of *cdrS* crRNA-spacer in the cell which directed the CRISPR known-down effect on *cdrS*. The forward primers used for PCR were located internal of *cdrS* crRNA-spacer, and the reverse primers were universal primers pUC13/M13 Rev located ~160 bp away from the downstream repeat motif in the mini-CRISPR array plasmid (Supplementary Figure 1). The PCR analysis showed the presence of *cdrS* crRNA-spacer in all the *cdrS* CRISPRi strains, and the tested corresponding complementation strains, suggesting that they did not escape CRISPRi-mediated editing via recombination between the repeat sequences flanking the crRNA-spacer on the plasmid. DNA ladder (right): 100 bp DNA Ladder (NEB); DNA ladder (left): MassRuler DNA Ladder Mix (Thermo Fisher Scientific).

### Supplementary Tables

**Supplementary Table 1. Genes up- and down regulated in *cdrS-ftsZ2* CRISPRi cells.**

From 3,595 genes transcripts were detected, altogether 97 genes had significant expression differences during *cdrS-ftZ2* repression, with 35 genes upregulated (Table S1A) and 62 genes downregulated (Table S1B), genes are sorted according to annotated functional category. The complete list of up- and downregulated genes can be found in the separate Supplementary Table 5 (Excel Table). All data with adjusted p-value p<0.005. TMD: predicted transmembrane domain.

**A. Upregulated genes**

The four upregulated genes HVO_B0193s2, HVO_B0193s, HVO_B0193 and HVO_B0192 (marked with an asterisk) might be part of an operon, HVO_B0192 and HVO_B0193 have already been shown to constitute an operon [1].

| Gene | Annotation | $\log_2$ |
|---|---|---|
| **Secreted, membrane, and cell surface protein** | | |
| HVO_1003 | GufA family transport protein (probable substrate zinc, signal peptide, TMD) | 2.0 |
| HVO_B0192* | conserved hypothetical protein (signal peptide, TMD) | 2.6 |
| HVO_2323 | conserved hypothetical protein (TMD) | 1.5 |
| HVO_0814 | chaperone (DnaJ domain, TMD) | 1.4 |
| HVO_2324 | pantothenate permease panF (TMD) | 1.2 |
| HVO_B0133 | conserved hypothetical protein (TMD) | 1.1 |
| HVO_0998 | conserved hypothetical protein (signal peptide, TMD) | 1.1 |
| HVO_B0132 | PQQ repeat protein (signal peptide, TMD) | 1.0 |
| HVO_0995 | conserved hypothetical protein (TMD) | 0.9 |
| HVO_2470 | sodium- and chloride-dependent transporter (TMD) | 0.8 |
| HVO_0883 | conserved hypothetical protein (signal peptide, TMD) | 0.7 |
| HVO_1869 | conserved hypothetical protein (TMD) | 0.7 |
| **Transport** | | |
| HVO_A0339 | ABC-type transport system periplasmic substrate-binding protein (probable substrate dipeptides/oligopeptides) | 1.0 |
| **Transcription** | | |
| HVO_B0319 | IclR family transcription regulator | 2.8 |
| HVO_B0193* | ArsR family transcription regulator | 2.9 |
| HVO_0576 | transcription regulator | 0.8 |
| HVO_A0394 | conserved hypothetical protein (ArsR-like helix-turn-helix domain) | 1.2 |
| **Signal transduction** | | |
| HVO_1358 | response regulator | 1.3 |
| HVO_1222 | CheR-like methyltransferase | 1.1 |
| **General metabolism** | | |
| HVO_A0470 | dioxygenase | 0.9 |
| HVO_2665 | HpcH/HpaI aldolase family protein | 0.7 |
| **Amino acid metabolism** | | |

| | | |
|---|---|---|
| HVO_2646 | dihydroxy-acid dehydratase (DHAD) | 1.1 |
| **DNA maintenance and repair** | | |
| HVO_1302 | DNA-directed DNA polymerase Y polY | 1.0 |
| HVO_A0450 | universal stress protein 3 | 0.9 |
| **Genes without known function** | | |
| HVO_A0129 | conserved hypothetical protein | 1.3 |
| HVO_B0193s2* | RNA of unknown function | 3.7 |
| HVO_B0193s* | RNA of unknown function | 3.5 |
| HVO_0582s | RNA of unknown function | 6.4 |
| HVO_2391s | RNA of unknown function | 2.3 |
| HVO_0259s | RNA of unknown function | 2.1 |
| HVO_2787s | RNA of unknown function | 1.1 |
| HVO_2021 | conserved hypothetical protein | 1.9 |
| HVO_2392 | conserved hypothetical protein | 1.8 |
| HVO_0457 | Zinc finger protein 330-like protein | 1.1 |
| HVO_B0195 | Hypothetical protein | 1.1 |

**B. Downregulated genes.**

| Gene | Annotation | $log_2$ |
|---|---|---|
| **secreted, membrane, and cell surface protein** | | |
| HVO_0739 | conserved hypothetical protein (TMD) | -3.3 |
| HVO_A0152 | conserved hypothetical protein (TMD) | -2.2 |
| HVO_A0493 | ABC-type transport system permease protein (probable substrate sugar, TMD) | -2.1 |
| HVO_A0173 | conserved hypothetical protein (TMD) | -1.9 |
| HVO_2034 | putative sugar ABC transporter permease (TMD) | -1.8 |
| HVO_1759 | putative iron-III ABC transporter permease (TMD) | -1.7 |
| HVO_2976 | carbon starvation protein CstA (TMD) | -1.6 |
| HVO_0607 | conserved hypothetical protein (Twin-arginine translocation pathway, signal sequence) | -1.6 |
| HVO_2064 | conserved hypothetical protein (signal peptide) | -1.6 |
| HVO_1228 | halocyanin (signal peptide) | -1.6 |
| HVO_A0318 | conserved hypothetical protein (TMD) | -1.3 |
| HVO_0343 | crcB protein-like protein (signal peptide, TMD) | -1.2 |
| HVO_A0165 | putative transporter (TMD) | -1.0 |
| HVO_1401 | putative sugar ABC transporter periplasmic substrate-binding protein (signal peptide) | -1.0 |
| HVO_1844 | conserved hypothetical protein (signal peptide) | -0.9 |
| HVO_B0063 | CbtB family protein (TMD) for cobalt transport | -2.4 |
| HVO_B0064 | CbtA family protein (signal peptide, TMD) for cobalt transport | -2.8 |
| HVO_B0108 | ABC-type transport system permease protein (TMD, Metl-like domain) | -2.3 |
| **Transport** | | |
| HVO_B0217 | ABC-type transport system periplasmic substrate-binding protein (probable substrate branched-chain amino acids) | -1.6 |
| HVO_A0494 | ABC-type transport system periplasmic substrate-binding protein (probable substrate sugar) | -1.8 |

| | | |
|---|---|---|
| **Cell division related** | | |
| HVO_0581 | cell division protein FtsZ2 | -2.6 |
| HVO_0392 | cell division protein SepF | -2.2 |
| HVO_0717 | cell division protein FtsZ1 | -1.3 |
| HVO_0689 | chromosome segregation protein SMC | -1.0 |
| **Transcription** | | |
| HVO_0582 | CdrS | -2.5 |
| HVO_0290 | ribbon-helix-helix CopG family protein | -1.6 |
| HVO_2110 | ArcR family transcription regulator | -1.4 |
| **Cobalamin (vitamin B12) biosynthesis** | | |
| HVO_B0054 | sirohydrochlorin cobaltochelatase | -2.1 |
| HVO_B0057 | cobalt-factor-III C17-methyltransferase | -2.6 |
| HVO_B0058 | cobalt-factor-III C17-methyltransferase | -2.1 |
| HVO_B0059 | cobalt-precorrin-5A hydrolase | -2.0 |
| HVO_B0060 | cobalt-precorrin-4 C11-methyltransferase | -2.8 |
| HVO_B0061 | cobalt-factor-II C20-methyltransferase | -2.7 |
| HVO_B0062 | precorrin-8W decarboxylase | -1.6 |
| HVO_A0488 | cob(I)alamin adenosyltransferase | -1.6 |
| HVO_0592 | adenosylcobinamide amidohydrolase | -1.5 |
| **General metabolism** | | |
| HVO_A0083 | Rieske-type [2Fe-2S] iron-sulfur protein | -2.0 |
| HVO_A0519 | monoamine oxidase regulatory protein | -1.8 |
| HVO_A0525 | enoyl-CoA hydratase | -1.8 |
| HVO_A0521 | phenylacetyl-coenzyme A ligase | -1.7 |
| HVO_0304 | electron transfer flavoprotein subunit alpha | -1.6 |
| HVO_2789 | putative molybdenum cofactor biosynthesis protein A | -1.4 |
| HVO_0585 | putative oxidoreductase | -1.0 |
| HVO_3014 | GTP-binding protein Era | -1.0 |
| HVO_1797 | mRNA 3´ end processing factor | -0.8 |
| HVO_0212 | putative lactoylglutathione lyase | -0.8 |
| HVO_B0238 | putative endoribonuclease L-PSP | -1.6 |
| HVO_1983 | malate synthase | -1.1 |
| HVO_B0200 | malate synthase | -0.9 |
| HVO_B0065 | thioredoxin-like superfamily protein | -2.2 |
| **Amino acid metabolism** | | |
| HVO_0041 | ornithine carbamoyltransferase (argF) | -1.4 |
| HVO_0043 | acetylornithine aminotransferase (argD) | -1.0 |
| HVO_0046 | rimK family protein | -1.0 |
| **Genes without known function** | | |
| HVO_B0055 | conserved hypothetical protein | -2.9 |
| HVO_3013 | conserved hypothetical protein | -0.8 |
| HVO_2973 | conserved hypothetical protein | -1.8 |
| HVO_B0240 | conserved hypothetical protein | -1.6 |
| HVO_2868s | RNA of unknown function | -1.4 |
| HVO_2073s | RNA of unknown function | -1.3 |
| HVO_2351s | RNA of unknown function | -1.2 |
| HVO_1106s | RNA of unknown function | -1.0 |
| HVO_1885s | RNA of unknown function | -2.2 |

**Supplementary Table 2. Proteins with significant changes in abundance.**

Proteins with significant changes in abundance from soluble (supernatant) and insoluble fractions sorted according to annotated functional category. The full list of quantified proteins can be found in the separate Supplementary Table 6 (Excel file). As a proportion of total protein-coding genes, an average of about 37% (1,499 proteins) proteome coverage was obtained.

Column On/Off and log$_2$: On, proteins are only found in CRISPRi cells; Off, proteins are only found in wild-type cells; log$_2$ ratio: log$_2$ values. SN: supernatant fraction; TMD: predicted transmembrane domain.

| | | wild-type vs CRISPRi differential abundance (On/Off and log$_2$ ratio) | |
|---|---|---|---|
| **Gene** | **Annotation** | **Pellet** | **SN** |
| **Secreted, membrane, and cell surface protein** | | | |
| HVO_B0153A | capsule biosynthesis CapC domain protein (signal peptide, TMD) | Off | |
| HVO_2492 | hypothetical protein (TMD) | Off | |
| HVO_A0629 | PAS domain, signal transduction histidine kinase domain (TMD) | Off | |
| HVO_0739 | hypothetical protein (TMD) | -4.96 | Off |
| HVO_2470 | sodium- and chloride-dependent transporter SNF | | -2.83 |
| HVO_1046 | hypothetical protein (TMD) | On | |
| HVO_2267 | hypothetical protein (TMD) | On | |
| HVO_2027 | DoxX domain protein (TMD) | | Off |
| **Transport** | | | |
| HVO_1110 | ABC-type transport system periplasmic substrate-binding protein (probable substrate cobalamin) | Off | |
| HVO_A0177 | ABC-type transport system ATP-binding protein | On | |
| HVO_1760 | Putative iron-III ABC transporter ATP-binding protein | | Off |
| HVO_2211 | TrkA family potassium uptake protein | On | |
| **Protein synthesis and translation** | | | |
| HVO_1148 | 30S ribosomal protein S15 | Off | |
| HVO_0809 | Met-tRNA synthetase | On | |
| HVO_0870 | Pro-tRNA synthetase | On | |
| HVO_1684 | Thr-tRNA synthetase | On | |
| HVO_0769 | TRAM domain-containing protein | On | |
| **Transcription** | | | |
| HVO_B0201 | IclR family transcription regulator | | Off |
| HVO_B0320 | IclR family transcription regulator | | Off |
| **Carbohydrate metabolism** | | | |
| HVO_1172 | galE UDP-glucose 4-epimerase | On | |
| HVO_1494 | fructose-1,6-bisphosphate aldolase | On | |
| HVO_0478 | glyceraldehyde-3-phosphate dehydrogenase type II | On | |
| HVO_1300 | triosephosphate isomerase | On | |
| HVO_2960 | Dihydrolipoyllysine-residue acetyltransferase | On | |
| HVO_B0085 | possible polygalacturonase, putative | | Off |
| **Central carbon metabolism** | | | |

| | | | |
|---|---|---|---|
| HVO_1000 | Acetyl-CoA synthetase | On | |
| **Amino acid metabolism** | | | |
| HVO_0044 | argB acetylglutamate kinase | On | |
| HVO_2852 | Succinylglutamate desuccinylase | | Off |
| HVO_2992 | phosphoribosyl-AMP cyclohydrolas | | Off |
| **Lipid metabolism** | | | |
| HVO_2725 | Isoprenyl diphosphate synthase (IdsA1) | On | |
| **General metabolism** | | | |
| HVO_0069 | arylsulfatase | On | |
| HVO_0869 | glutamate synthase subunit | On | |
| HVO_0884 | Aldehyde reductase | On | |
| HVO_1874 | probable oxidoreductase (aldo-keto reductase family protein) | On | |
| HVO_0662 | ThiN homolog with a predicted N-terminal helix-turn-helix (HTH) DNA binding domain | On | |
| HVO_1009 | Oxidoreductase related to aryl-alcohol dehydrogenases | On | |
| HVO_2336 | pyridoxal 5'-phosphate synthase lyase subunit PdxS) | On | |
| HVO_2348 | GTP cyclohydrolase I | On | |
| HVO_2650 | 4-hydroxybenzoate 3-monooxygenase | On | |
| HVO_2790 | ATP-binding protein Mrp | On | |
| HVO_1697 | FAD-dependent oxidoreductase (GlcD/DLD_GlcF/GlpC domain fusion protein) | | 2.79 |
| HVO_2579 | nicotinate-nucleotide pyrophosphorylase (carboxylating) | | -2.35 |
| HVO_0510 | dihydroneopterin aldolase, archaeal-type, MptD | | -2.15 |
| HVO_2127 | M20 family amidohydrolase (homolog to indole-3-acetyl-aspartate hydrolase) | | Off |
| HVO_2848 | probable PrkA-type serine/threonine protein kinase, PrkA1 | | Off |
| **DNA maintenance and repair** | | | |
| HVO_2452 | ribonucleoside-diphosphate reductase, adenosylcobalamin-dependent | On | |
| HVO_0203 | Replication factor C small subunit | On | |
| HVO_0104 | DNA repair and recombination protein RadA | 3.84 | |
| HVO_2911 | deoxyribodipyrimidine photo-lyase, phr1 | | Off |
| **Cell division-related** | | | |
| HVO_0392 | SepF | | -3.32 |
| HVO_0581 | FtsZ2 | | -3.19 |
| **Proteins without known function** | | | |
| HVO_2519 | hypothetical protein | On | |
| HVO_0400 | hypothetical protein | On | |
| HVO_1535 | hypothetical protein | | Off |
| HVO_0508 | hypothetical protein | On | |
| HVO_1377 | uncharacterised protein family UPF0145 | On | |
| HVO_0577 | uncharacterised DUF2028 domain | | -4.55 |

**Supplementary Table 3. Quantitative analysis of cell shape**

The cell division mutants were quantified as four major distinct cell shapes: filaments (Cell area >= 6.5 µm$^2$ , Circularity < =0.4), giant plate cells (Cell area >= 6.5 µm$^2$ , Circularity > 0.4); wild-type-like cells (cell area between 2 µm$^2$ and 6.5 µm$^2$ ); cellular debris (cell area < 2 µm$^2$).
Analysed were WT (WT x pTA232), *cdrS* repression (*cdrS* CRISPRi#1-3) (Fig. 5A) and overexpression (WT x *cdrS*) (Fig. 7D) cells.

| | WT X pTA232 (n=406) | *cdrS* CRISPRi#1 (n=314) | *cdrS* CRISPRi#2 (n=330) | *cdrS* CRISPRi#3 (n=173) | WT x *cdrS* (n=131) |
|---|---|---|---|---|---|
| Filaments | 0 % | 20 % | 30 % | 52.6 % | 2.3 % |
| Giant cells | 0 % | 5.7 % | 3 % | 4.6 % | 45 % |
| Wild-type like | 100 % | 43 % | 52.4 % | 24.9% | 40.5 % |
| Cellular debris | 0 % | 31.2 % | 14.5 % | 18 % | 12.2 % |

**Supplementary Table 4A. Strains used**

| **Strains** | **Genotype** | **Reference** |
|---|---|---|
| H26 | ΔpHV2, Δ*pyrE2* | (Allers, Ngo, Mevarech, & Lloyd, 2004) |
| HV30 | ΔpHV2, Δ*pyr*E2, Δ*trp*A, Δ*leu*B, Δ*cas*6, Δ*cas*3, Δ*bga*H | (Stachler & Marchfelder, 2016) |
| HV32 | ΔpHV2, Δ*pyrE2*, Δ*trpA*, Δ*leuB, Δl*-B, ΔHVO_2.385.045–2.386.660, ΔHVO_pHV4: 204.834-218.566 | (Stachler & Marchfelder, 2016) |
| HV35 | ΔpHV2, Δ*pyrE2*; Δ*leuB*; Δ*trpA*; ΔHVO_2,385,045–2,386,660::p.tna,*cas6*,*cas8*, *cas7*,*cas5*, t.syn; ΔHVO_pHV4:204,834-218,566 | this study |
| DH5α | F- 80*lac*ZΔM15 Δ(*lac*ZYA-*arg*F) U169 *rec*A1 *end*A1 *hsd*R17 (rk-, mk+) *gal*- *pho*A *sup*E44λ- *thi*-1 *gyr*A96 *rel*A1 | Invitrogen (Thermo Fisher Scientific, Waltham, USA) |
| GM121 | F- *dam-3 dcm-6 ara-14 fhuA31 galK2 galT22 hdsR3 lacY1 leu-6 thi-1 thr-1 tsx-78* | (Allers, Barak, Liddell, Wardell, & Mevarech, 2010) |

**Supplementary Table 4B. Plasmids used**

| plasmids | relevant properties | Reference/ source |
|---|---|---|
| pBlueScriptII | *E. coli* plasmid | Stratagene |
| pBlue-HVO_0582 | *E. coli* plasmid with HVO_0582 *gene* | this study |
| pTA232 | Shuttle vector with *leu*B marker and pHV2 replication origin | (Allers et al., 2004) |
| pTA231 | Shuttle vector with *trpA* marker, pHV2 replication origin | (Allers et al., 2004) |
| pTA409 | Shuttle vector with *pyrE*2 marker and pHV1 replication origin | (Delmas, Shunburne, Ngo, & Allers, 2009) |
| pTA131-up-HVO_0582-do | ColE1 ori, f1 ori*, lac*Z*,* AmpR*, pyr*E2, gene HVO_0582 with 500 bp flanking upstream and downstream regions | this study |
| pTA131-up-ΔHVO_0582-do | ColE1 ori, f1 ori*, lac*Z*,* AmpR*, pyr*E2, 500 bp flanking regions upstream and downstream of HVO_0582 | this study |
| pTA131-Cup-p.ntaA-cas6b8b75-t.Syn-Cdo | ColE1 ori, f1 ori, *lacZ*, AmpR, *pyrE*2, Cup-p.ntaA*cas6bcas8bcas7cas5*-t.syn-Cdo | this study |
| pMA-telecrRNA | *E. coli* plasmid with promoter, crRNA against spacer C1, flanked by t-Elements, terminator | (Maier, Dyall-Smith, & Marchfelder, 2015) |
| pTA927-ptnaA-NFlag | Shuttle vector with *pyrE2* marker, pHV2 replication origin, a tryptophan inducible promoter and a 3xFLAG tag cDNA | (Fischer et al., 2010) |
| pMK-RQ-0582anti#1 | *E. coli* plasmid containing the promoter, spacer sequence flanked by Haloferax repeats and terminator, expressing a crRNA#1 against the HVO_0582 RNA gene | GeneArt (Thermo Fisher Scientific) |
| pMK-RQ-0582anti#2 | *E. coli* plasmid containing the promoter, spacer sequence flanked by Haloferax repeats and terminator, expressing a crRNA#2 against the HVO_0582 RNA gene | GeneArt (Thermo Fisher Scientific) |
| pMK-RQ-0582anti#3 | *E. coli* plasmid containing the promoter, spacer sequence flanked by Haloferax repeats and terminator, expressing a crRNA#3 against the HVO_0582 RNA gene | GeneArt (Thermo Fisher Scientific) |
| pTA232-0582anti#1 | Plasmid containing the promoter, spacer sequence flanked by *Haloferax* repeats and terminator, expressing a crRNA#1 against the template strand of the HVO_0582 gene | this study |
| pTA232-0582anti#2 | Plasmid containing the promoter, spacer sequence flanked by *Haloferax* repeats and | this study |

| | | |
|---|---|---|
| | terminator, expressing a crRNA#2 against the template strand of the HVO_0582 gene | |
| pTA232-0582anti#3 | Plasmid containing the promoter, spacer sequence flanked by *Haloferax* repeats and terminator, expressing a crRNA#3 against the template strand of the HVO_0582 gene | this study |
| pMA-tele-anti#1 | *E. coli* plasmid with promoter, crRNA anti#1 against HVO_0582 , flanked by t-elements, terminator | this study |
| pMA-tele-anti#2 | *E. coli* plasmid with promoter, crRNA anti#2 against HVO_0582 , flanked by t-elements, terminator | this study |
| pMA-tele-anti#3 | *E. coli* plasmid with promoter, crRNA anti#3 against HVO_0582 , flanked by t-elements, terminator | this study |
| pTA232-tele-anti#1 | pTA232 plasmid with promotor, crRNA anti#1 against HVO_0582, flanked by t-elements, terminator | this study |
| pTA232-tele-anti#2 | pTA232 plasmid with promotor, crRNA anti#2 against HVO_0582, flanked by t-elements, terminator | this study |
| pTA232-tele-anti#3 | pTA232 plasmid with promotor, crRNA anti#3 against HVO_0582, flanked by t-elements, terminator | this study |
| pTA927-ptnaA-HVO_0582NFlag | Plasmid pTA927 with tryptophan inducible promoter, HVO_0582 fused to an N-terminal 3xFlag tag | this study |
| pTA231-pfdx-HVO_0582NFlag | Plasmid pTA231 with HVO_0582 fused with an N-terminal Tag under the control of the pfdx promoter | this study |
| pTA409-pfdx-HVO_0582-nat.t | Plasmid expressing the gene HVO_0582 under the control of the pfdx promotor and the natural terminator | this study |
| pTA409-pfdx-HVO_0581-nat.t | Plasmid expressing the gene HVO_0581 under the control of the pfdx promotor and the natural terminator | this study |
| pTA409-pfdx-HVO_0582-HVO_0582-nat.t. | Plasmid expressing the operon HVO_0582-HVO_0581 under the control of the pfdx promotor and the natural terminator | this study |

**Supplementary Table 4C. Oligos used**

| primer | 5’->3’ sequence |
|---|---|
| RS | CACAGGAAACAGCTATGACC |
| US | GTAACGCCAGGGTTTTCCC |
| P1upfw | GCGTCGGCTCGATTCCACTCACCAACG |
| P2dorev | CCTCGACGCCGTCGAGAGACTCGAATC |
| anti#2 fw | (Phos)GCGTAAGACGGTTGTCTTGTTTCAGACGAACCCTTGTGG |
| anti#2 rv | (Phos)GATAATCACCACAGAGGGGCTTCAACTACCGATCAACG |
| anit#3 fw | (Phos)ACCCGACGGCGGCCCGTAGTTTCAGACGAACCCTTGTGG |
| anti#3 rv | (Phos)AGCTGTAAGACAACCGTCGCTTCAACTACCGATCAACG |
| 5’-HVO_0582-*Hind*III | AAGCTTATGGAGCGTGTGACACTACGAATTCC |
| 3’-HVO_0582-*Xba*I | TATATCTAGATTACACCTTTGCCCAGCCGCG |
| 5‘-HVO_0582-*Nde*I | TATTACATATGGAGCGTGTGACACTACGAATTCC |
| 3'-HVO_0582-*Hind*III | TATATAAAGCTTTTACACCTTTGCCCAGCC |
| 5‘-HVO_0581-*Nde*I | TATATACATATGATGCAGGATATCGTTCGC |
| 3'-HVO_0581-nat.t.-*Apa*I | TATATAGGGCCCCTCGCGGTCGAAGAAATC |
| 5‘-*Hind*III-nat.t | TATATAAAGCTTCGCCCTGTCCGACCCGCG |
| 5-HindIII-Cas8 | TATTATAAGCTTACAGGTCCAGATATCGACGACTTCG |
| 8R126A#2 | CCACGAACGAACGGCTCCGAGAATCGTGCTGGC |
| CdelupKpnI | TATAGGTACCCGCTCGTCGGTGAGTCGCTCACCGACTTCCG |
| CdelupiEcoRV | TATAGATATCCGAGGCGGAGCGTCGAGAGCGCTAGTC |
| 3'-HVO_B0192-NB-rev | GCGAGCGAGACGATAAGCCAGTCG |
| 5'-HVO_B0193-NB-fw | CAACCATTACAGCGTCTACACCGC |
| 5'-HVO_0739-NB-fw | CGGGGGGTTCCTCCTCGGCGTGC |
| 3'-HVO_0739-NB-rev | GAAGAGCGCGACGCCGAGGACGGC |
| anti#1 fw | (Phos)GATTATCGCGTAAGACGGGTTTCAGACGAACCCTTGTGG |
| anti#2 rev | (Phos)ACCACAGAGGGATAGAATGCTTCAACTACCGATCAACG |
| HVO_0582-UP | TCGAACGGCGAAT CTCGCGTAAACGC |
| HVO_0582-DO | CCTCGGCCTTGAC CGTGCGGGCGC |
| iPCR_HVO_0582-Do | [P]CAGCAAGCGCG GCTGGGCAAAG |
| iPCR_HVO_0582-UP | [P]GTTTACATTCCC CCGGTAAGACGG |
